# Liver-specific suppression of ANGPTL4 improves obesity-associated diabetes and mitigates atherosclerosis in mice

**DOI:** 10.1101/2020.06.02.130922

**Authors:** Abhishek K. Singh, Balkrishna Chaube, Alberto Canfrán-Duque, Xinbo Zhang, Nathan L. Price, Binod Aryal, Jonathan Sun, Kathryn M Citrin, Noemi Rotllan, Richard G. Lee, Yajaira Suárez, Carlos Fernández-Hernando

## Abstract

Angiopoietin-like 4 (ANGPTL4) is a major regulator of lipoprotein lipase (LPL) activity, which is responsible for maintaining optimal levels of circulating triacylglycerol (TAG) for distribution to different tissues including the adipose tissues (ATs), heart, muscle and liver. Dysregulation of trafficking and portioning of fatty acids (FA) can promote ectopic lipid accumulation in metabolic tissues such as the liver, ultimately leading to systemic metabolic dysfunction. To investigate how ANGPTL4 regulates hepatic lipid and glucose metabolism, we generated liver-specific ANGPTL4 knockout mice (*LKO*). Using metabolic turnover studies, we demonstrate that hepatic ANGPTL4 deficiency facilitates catabolism of TAG-rich lipoprotein (TRL) remnants in the liver via increased hepatic lipase (HL) activity, which results in a significant reduction in circulating TAG and cholesterol levels. Deletion of hepatocyte ANGPTL4 protects against diet-induce obesity, glucose intolerance, liver steatosis, and atherogenesis. Mechanistically, we demonstrate that absence of ANGPTL4 in hepatocytes promotes FA uptake which results in increased FA oxidation, ROS production, and AMPK activation. Finally, we demonstrate the utility of a targeted pharmacologic therapy that specifically inhibits ANGPTL4 in the liver and protects against diet-induced obesity, dyslipidemia, glucose intolerance, and liver damage without causing any of the deleterious effects previously observed with neutralizing antibodies.

## INTRODUCTION

The liver is a central metabolic organ that is essential for regulating systemic glucose and lipid homeostasis (1, 2). Lipid homeostasis is maintained by the liver via a regulated balance between lipid acquisition (FA uptake and biosynthesis) and removal (FA oxidation (FAO) and VLDL secretion) (3-5). Dysregulation in these critical processes leads to the accumulation of excess TAG in hepatocytes, resulting in the development of non-alcoholic fatty liver disease (NAFLD) and associated disorders such as hepatic insulin resistance (IR), type 2 diabetes (T2D) and atherosclerosis (5-10). The key pathological feature of NAFLD is the accumulation of intra-hepatic triacylglycerol (TAG), owing to increased flux of free FA (FFA) derived from lipolysis in adipose tissue (AT), dietary chylomicrons, and intrahepatic de novo lipogenesis (DNL) (5, 11). Lipoprotein lipase (LPL) plays a vital role in the homeostasis of lipid metabolism at the systemic level. However, the cellular mechanisms behind the regulation of lipid homeostasis in the liver remain unclear and are now a central focus in the field.

ANGPTL4 is a multifaceted secreted protein that is highly expressed in metabolic tissues, most prominently in adipose tissue (AT) and liver (12-14). ANGPTL4 regulates many cellular and physiological functions, mainly via inhibiting LPL activity at the posttranslational level (15-20). Human genetics and clinical studies have emphatically demonstrated that mutations in the ANGPTL4 gene (E40K) are associated with reduced plasma TAG and glucose levels (21-24). Importantly, the circulating level of ANGPTL4 is positively correlated with increased risk of cardiovascular disease (CVD) and T2D, along with obesity-associated diabetic phenotypes such as high BMI, fat mass, and altered glucose homeostasis (25). These observations indicate that ANGPTL4 is strongly implicated in the development of metabolic diseases in humans; however, the underlying molecular mechanism is poorly understood. Early work on ANGPTL4 in rodents was performed using whole-body transgenic or knockout mouse models that limit our ability to understand the specific role of ANGPTL4 in vital metabolic tissues. Therefore, most of the conclusions drawn from these studies have been contradictory (13, 21, 26-30). Multiple groups have shown that loss of ANGPTL4 using knockout models or monoclonal antibodies against ANGPTL4 results in severe gut inflammation upon high-fat diet (HFD) or western type diet (WD) feeding, leading to metabolic complications and reduced survival (26, 31). In these studies, it is not clear whether these effects of ANGPTL4 deficiency are LPL dependent or independent. Therefore, future research on ANGPTL4 must be conducted in mouse models that are not affected by these confounding factors. To address this significant gap in our understanding of how ANGPTL4 functions, we have generated novel mouse models that lack expression of ANGPTL4 specifically in the adipose tissue or liver, where ANGPTL4 is predominantly expressed.

Our recent work has demonstrated an essential role for adipose-derived ANGPTL4 in regulating lipid uptake and regulation of metabolic function by AT (32). We have observed that adipose-specific ANGPTL4 knockout (Ad-KO) mice displayed reduced circulating TAG levels, similar to what was observed in humans with a missense mutation in ANGPTL4 (E40K variant). However, other factors identified in this study such as body weight and regulation of glucose homeostasis, were unaltered after chronic administration of HFD feeding (32). These findings suggest that ANGPTL4 in other tissues, such as the liver, could also play a key role in the regulation of whole-body metabolism. Moreover, the physiological roles of ANGPTL4 in liver metabolism are primarily unclear. Therefore, we generated a novel liver-specific ANGPTL4 knockout mouse model (*LKO*) to investigate the contribution of hepatic ANGPTL4 in obesity-linked metabolic disorders. We demonstrate that hepatic depletion of ANGPTL4 improves metabolic parameters such as body weight, serum lipid profile, ectopic lipid accumulation, and insulin sensitivity, and attenuates atherogenesis. Notably, we uncover a novel cellular mechanism for regulation of lipid homeostasis downstream of ANGPTL4 involving hepatic lipase (HL) and redox-dependent (ROS) activation of AMP-activated protein kinase (AMPK). Lastly, we evaluated the therapeutic potential of ANGPTL4 for the treatment of obesity-linked diabetes through liver-specific silencing of ANGPTL4 with GalNac-conjugated antisense oligonucleotides (ANGPTL4-ASO) in the mouse, as a preclinical study.

## RESULTS

### Hepatic deletion of ANGPTL4 alters systemic lipid metabolism

To determine the tissue-specific function of ANGPTL4, we first assessed its tissue distribution. We analyzed RNA sequencing data from GTEx (https://gtexportal.org/home/). This data shows that ANGPTL4 is highly expressed in the human liver. (**Figure 1A**). To understand the liver-specific role of ANGPTL4 on whole-body lipid and glucose metabolism, we generated and *Angptl4* conditional knockout mice, which were then bred with Albumin-Cre animals to specifically delete the expression of *Angptl4* in the liver (*LKO*). *Angptl4* mRNA expression analyzed by qPCR was reduced by more than 95% in the liver of *LKO* mice as compared to WT mice (**Figure 1B** and **C**), with no compensatory increase in the expression of either ANGPTL3 or ANGPTL8 (**Figure S1**). To assess whether hepatic ANGPTL4 deficiency influences lipoprotein and glucose metabolism, we measured body weight and plasma levels of lipids and glucose in 2-month-old *LKO* mice fed a chow diet (CD). While fasting blood glucose levels and body weight were unaltered, we observed significantly lower circulating TAGs, total cholesterol (TC), and HDL-C in *LKO* mice (**Figure 1D-F**). Moreover, we found a reduction in the levels of TAGs and cholesterol in the VLDL and HDL fractions of *LKO* mice, respectively (**Figure 1G**). These findings indicate that hepatic ANGPTL4 controls circulating lipid levels.

**Figure 1.**
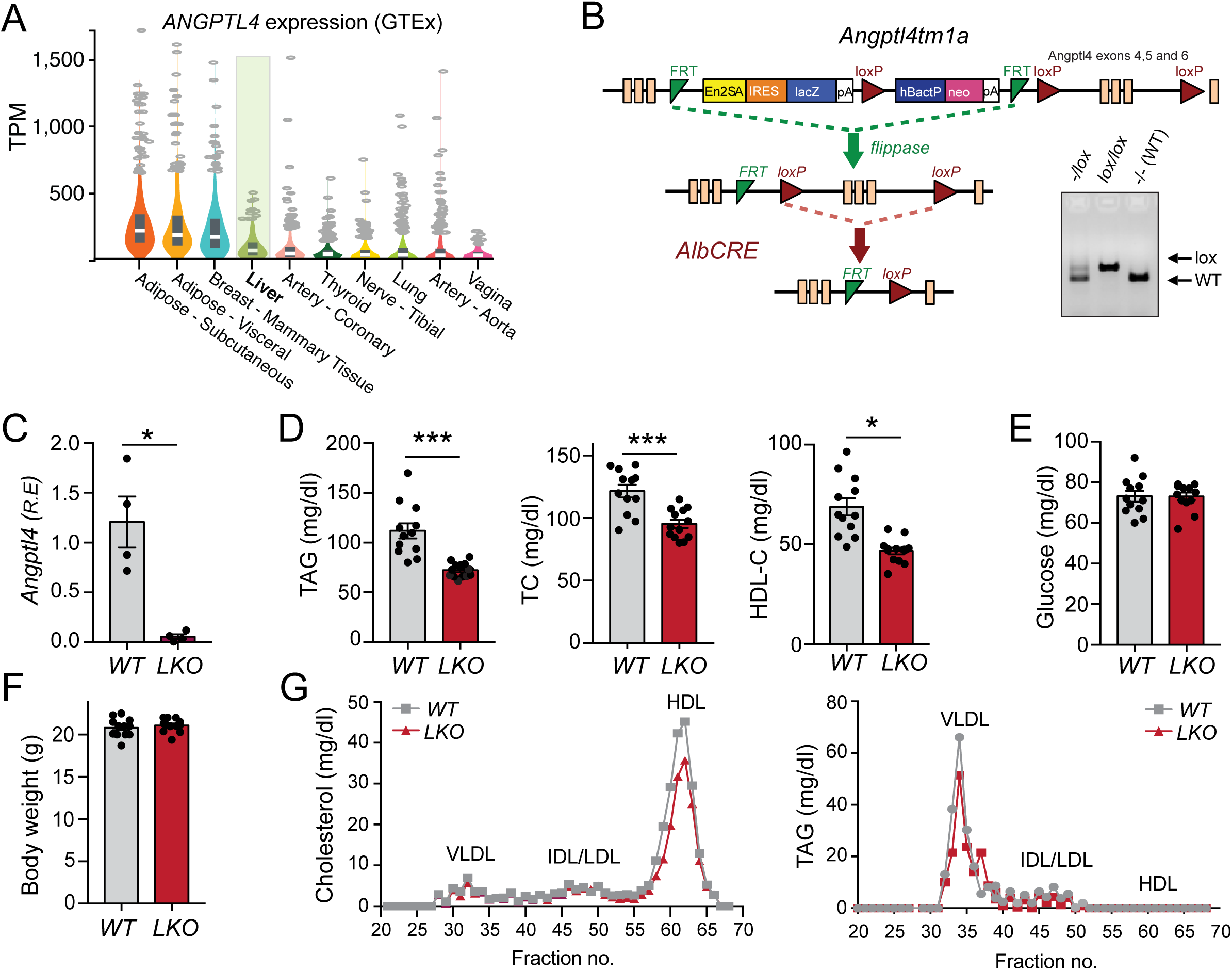
The loss of ANGPTL4 in the liver improves the serum lipid profile. (**A**) Analysis of human ANGPTL4 expression in different tissues using GTEx database. Liver is highlighted. TPM represents transcript per millions reads. (**B**) Schematic diagram illustrating the generation of liver-specific ANGPTL4 knockout (*LKO*) mice. The construct is composed of a short flippase recombination enzyme (Flp)-recognition target (FRT), reporter, and a Cre recombinase recognition target (loxP). Angptl4 exons 4-6 are flanked by lox P site. Mice with the floxed allele were generated by crossing with flp recombinase-deleter mice (mid panel). Subsequently, these mice were bred with mice expressing Cre recombinase to produce tissue specific ANGPTL4 knockout mice (bottom). PCR amplification of *Angptl4*^*fl/fl*^ mice displaying bands from both, one, or none of the alleles floxed (Right panel). (**C**) mRNA expression of *Angptl4* in the liver of WT & *LKO* mice. (**D**) Fasted plasma triacylglycerol (TAG; left panel), total cholesterol (middle panel), and HDL-C (right panel) from WT and *LKO* mice chow diet (CD) for two months (n=10-13). (**E** and **F**) Blood glucose levels (**E**), and body weight (**F**) of WT and *LKO* mice fed a chow diet (CD) for two months (n=10-13). (**G**) Cholesterol (left panel) and TAG (right panel) content of FPLC-fractionated lipoproteins from pooled plasma of WT and *LKO* mice fed a CD for two months (n=7). R.E denotes relative expression. All data represent mean ± SEM. *p<0.05, ***p<0.001 comparing *LKO* with WT mice using the unpaired t-test.

### Absence of ANGPTL4 in the liver enhances hepatic lipid uptake

In order to understand how liver-derived ANGPTL4 regulates systemic lipid metabolism, we assessed possible causes of hypotriglyceridemia in *LKO* mice. We reasoned that these effects could be due to either enhanced Triglyceride-rich lipoprotein (TRL) (chylomicrons and VLDL) catabolism, reduction in hepatic VLDL production, or defective lipid absorption in the gut. To determine the impact of hepatocyte-specific suppression of ANGPTL4 on the clearance of dietary TAGs (chylomicrons) in circulation, we administered mice with an intra-gastric gavage of olive oil and plasma TAGs and FFAs were measured at indicated time points post-gavage. We noticed reduced levels of postprandial plasma TAGs and FFAs in *LKO* mice as compared to WT mice (**Figure 2A** and **B**). Further, to identify the tissue(s) responsible for the enhanced lipid clearance, we analyzed lipid uptake using a [^3^H]-oleate-labeled triolein tracer. *LKO* mice displayed increased lipid clearance by the liver as compared to the WT littermates (**Figure 2C**). Next, we determined whether the accelerated clearance of TRL particles was due to an increase in the lipolytic activity of plasma. Therefore, we analyzed the activity of the two crucial enzymes involved in the hydrolysis of TAG, LPL and HL, in the plasma after heparin injection. Remarkably, *LKO* mice had markedly increased post-heparin plasma HL activity as compared to WT mice (**Figure 2D**). We also noticed a modest increase in post-heparin plasma LPL activity in *LKO* mice (**Figure 2E**).

**Figure 2.**
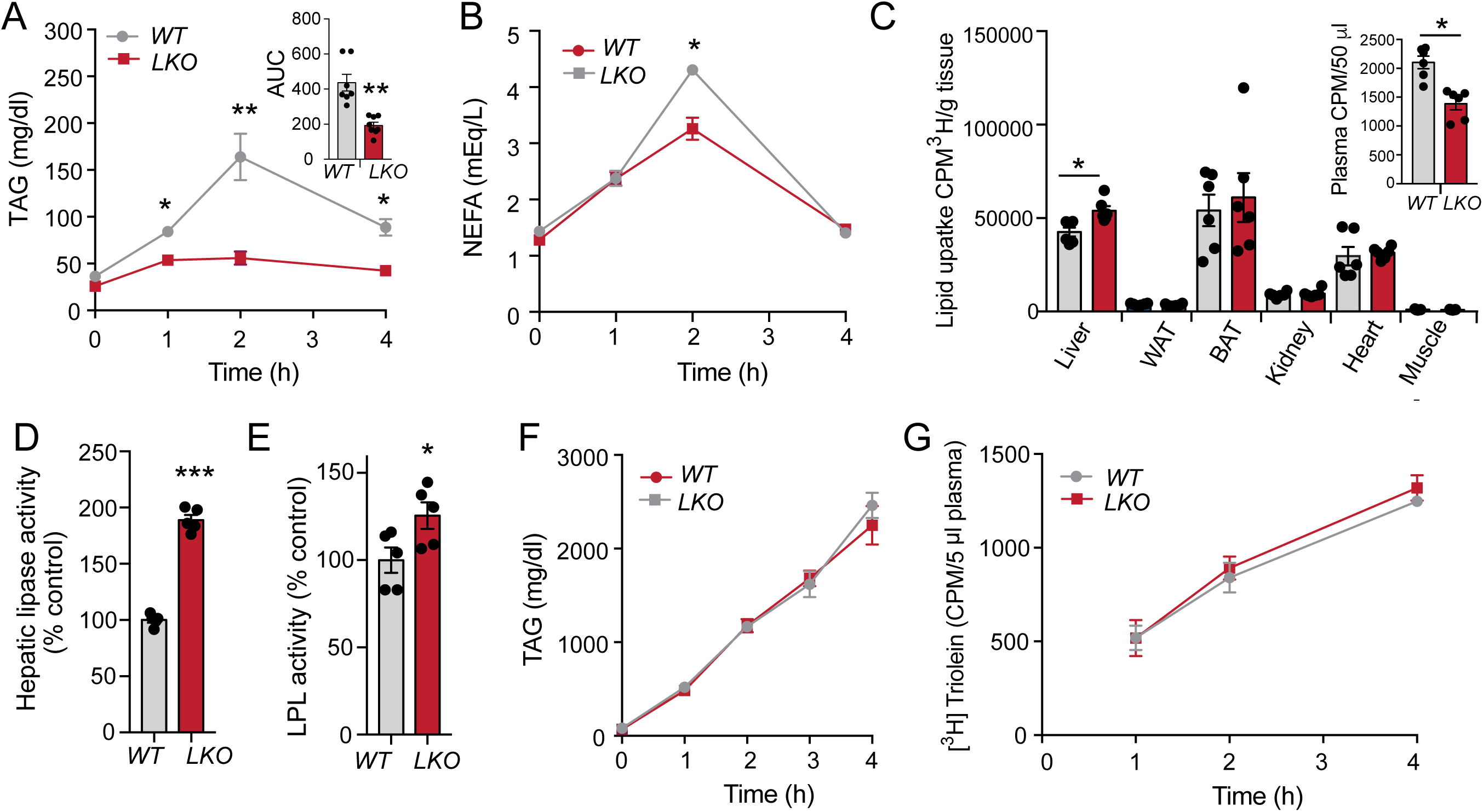
Absence of ANGPTL4 in the liver enhances plasma TAG clearance and hepatic lipid uptake. (**A** and **B**) Oral lipid tolerance test showing the clearance of TAG and NEFA from the plasma of WT and *LKO* mice fasted for 4h followed by oral gavage of olive oil. Inset represent the AUC. (**C**) Radioactivity incorporation in indicated tissues after 2h of oral gavage of [^3^H]-labeled triolein in WT or *LKO* mice fasted for 4h. Inset represent plasma lipid clearance. (**D** and **E**) HL and LPL activity in the post-heparin plasma from WT and *LKO* mice. (**F**) Plasma TAG from overnight fasted WT and *LKO* mice treated with plasma LPL inhibitor poloxamer 407 to inhibit the hydrolysis of circulating TAG (n=5). (**G**) Serum ^3^H counts after injection of poloxamer 407 combined with oral lipid gavage containing ^3^H-triolein (n=5). All data represent the mean ± SEM. *p<0.05 **p<0.01 ***p<0.001 comparing *LKO* with WT mice using the unpaired t-test.

To determine whether hepatic VLDL secretion is also altered in response to depletion of ANGPTL4 in the liver, we used poloxamer 407, which prevents uptake of plasma lipids by inhibiting LPL and HL mediated TAG hydrolysis. We did not find any difference in the VLDL TAG secretion between the groups (**Figure 2F**). Next, we analyzed lipid absorption by the intestine. Mice were injected intraperitoneally with poloxamer 407 and administrated an oral gavage with [^3^H]-oleate-labeled triolein. However, no significant changes were observed in the appearance of ^3^H-triolein in the plasma of *LKO* mice as compared to WT (**Figure 2G**). These observations suggest that differences in plasma lipids are not related to changes in either hepatic lipoprotein production or intestinal lipid absorption. These data indicate that hepatic deficiency of ANGPTL4 reduces circulating TAGs by enhancing lipolysis and clearance of TRL in the liver.

### Lack of ANGPTL4 in the liver reduces body weight, fat mass, and intrahepatic lipids

Since elevated circulating TAGs induce ectopic lipid deposition during obesity-induced insulin resistance (5, 7, 10, 33, 34) and mice lacking ANGPTL4 in the liver showed lower plasma TAG levels, we tested whether liver-specific depletion of ANGPTL4 could protect against elevated metabolic stress during high-fat diet (HFD) feeding. *LKO* and WT mice were challenged with a HFD (60% kcal from fat) for 16 weeks. Body weight and food intake were monitored weekly, and at the end of the study, metabolic parameters were measured. We observed that *LKO* mice showed a significant reduction in body weight as compared to WT littermates over 16 weeks on HFD (**Figure 3A**; **Figure S2A**), while no difference in food intake was observed between the groups (**Figure S2B**). This was accompanied by lower total body fat mass, reduced fat weight, and reduced cell size of adipocytes in the *LKO* mice (**Figure 3B-D**; **Figure S2C**).

**Figure 3.**
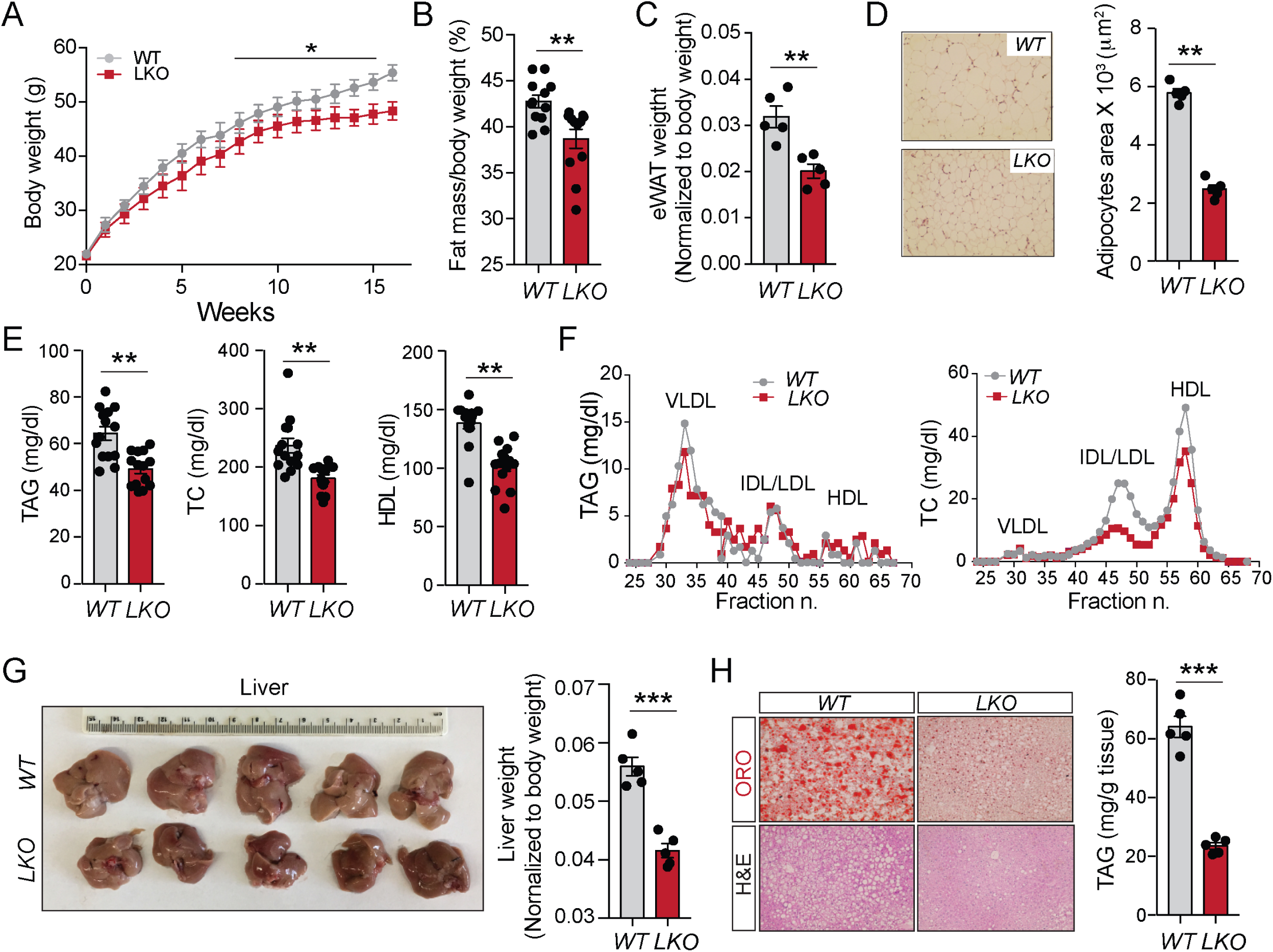
Hepatic ANGPTL4 deficiency protects from diet-induced obesity and decreases hepatic lipid accumulation. (**A** and **B**) Body weight and fat mass measured by Echo-MRI of WT and *LKO* mice fed a high–fat diet (HFD) for 16 weeks. (n=10-13). (**C** and **D**) Fat (eWAT) weight and representative images of H&E sections of WAT and quantification of adipocyte size isolated from WT and *LKO* mice fed an HFD for 16 weeks. (**E**) Levels of TAG, total cholesterol (TC), and HDL-C in the plasma of WT and *LKO* mice fed an HFD for 16 weeks. (**F**) FPLC analysis of lipoprotein profile from pooled plasma of WT and *LKO* mice (n=5). (**G** and **H**) Representative images of the liver, Liver weight, hepatic TAG levels, and representative pictures of Oil Red O and H&E-stained sections of liver isolated from WT and *LKO* mice fed HFD for 16 weeks. All data represent the mean ± SEM. *p<0.05 **p<0.01 ***p<0.001 comparing *LKO* with WT mice using the unpaired t-test.

Moreover, similar to the findings observed in CD-fed mice, we also found a significant reduction in circulating TAGs, TC, and HDL-C in *LKO* mice fed a HFD (**Figure 3E**). These results were further confirmed by lipoprotein fractions analysis (**Figure 3F**). Since alterations in body fat mass are often associated with changes in lipid accumulation in the liver (35), we also analyzed neutral lipid content in the liver of *LKO* and WT mice. As anticipated, we noticed a significant decrease in the liver weight of *LKO* mice as compared to WT mice (**Figure 3G**). Oil-Red O and H&E staining of liver sections further indicated reduced accumulation of neutral lipids in the liver of *LKO* mice (**Figure 3H**, left panels). Furthermore, we also found a marked reduction in hepatic TAG levels in *LKO* mice (**Figure 3H**, right panel). These data demonstrate that genetic deficiency of *Angptl4* in the liver protects against hepatic steatosis.

Elevated lipid deposition in the liver promotes the induction of hepatic inflammation (11), therefore we investigated whether loss of ANGPTL4 in the liver influences hepatic inflammation under HFD fed conditions. We analyzed the total number of monocytes and Kupffer cells (KC) in the liver of mice challenged with HFD for 16 weeks. FACS analysis showed that lack of hepatic ANGPTL4 reduced the number of monocytes in the liver, though we did not observe change in the number of KC (**Figure S2D**). In a nutshell, these data indicate that hepatic ANGPTL4 controls lipoprotein metabolism, and its absence improves the metabolic health of mice during pathophysiological conditions such as diet-induced obesity.

### Ablation of hepatic ANGPTL4 improves glucose homeostasis and enhances insulin sensitivity in metabolic tissues

Recent studies on humans and mice have reported that loss of function of ANGPTL4 is associated with improved glucose tolerance and insulin sensitivity (21, 27). Since diet-induced obesity is often accompanied by insulin resistance-linked glucose intolerance, we assessed the net functional outcome of hepatic ANGPTL4 deficiency on systemic glucose metabolism during diet-induced obesity. We found that *LKO* mice showed remarkably improved glucose tolerance as compared to WT after 16 weeks of HFD feeding (**Figure 4A**). One possible explanation for the improved glucose homeostasis in the *LKO* mice is improved insulin sensitivity. To test this hypothesis, we performed insulin tolerance tests (ITT) on *LKO* and WT mice fed with HFD for 16 weeks. Insulin-stimulated glucose disposal was significantly increased in *LKO* mice as compared to WT controls (**Figure 4B**). To independently evaluate insulin sensitivity in different metabolic tissue(s), we assessed the levels of phosphorylated AKT following insulin treatment, a well-established *in vivo* biochemical approach to measure insulin sensitivity. p-AKT/AKT ratio was markedly enhanced in metabolic tissues such as muscle, adipose tissue, and liver of *LKO* mice following intraperitoneal injection of insulin (**Figure 4C**). Overall, these results demonstrate that genetic deficiency of ANGPTL4 in the liver improves glucose homeostasis and augments systemic insulin sensitivity during obesity.

**Figure 4.**
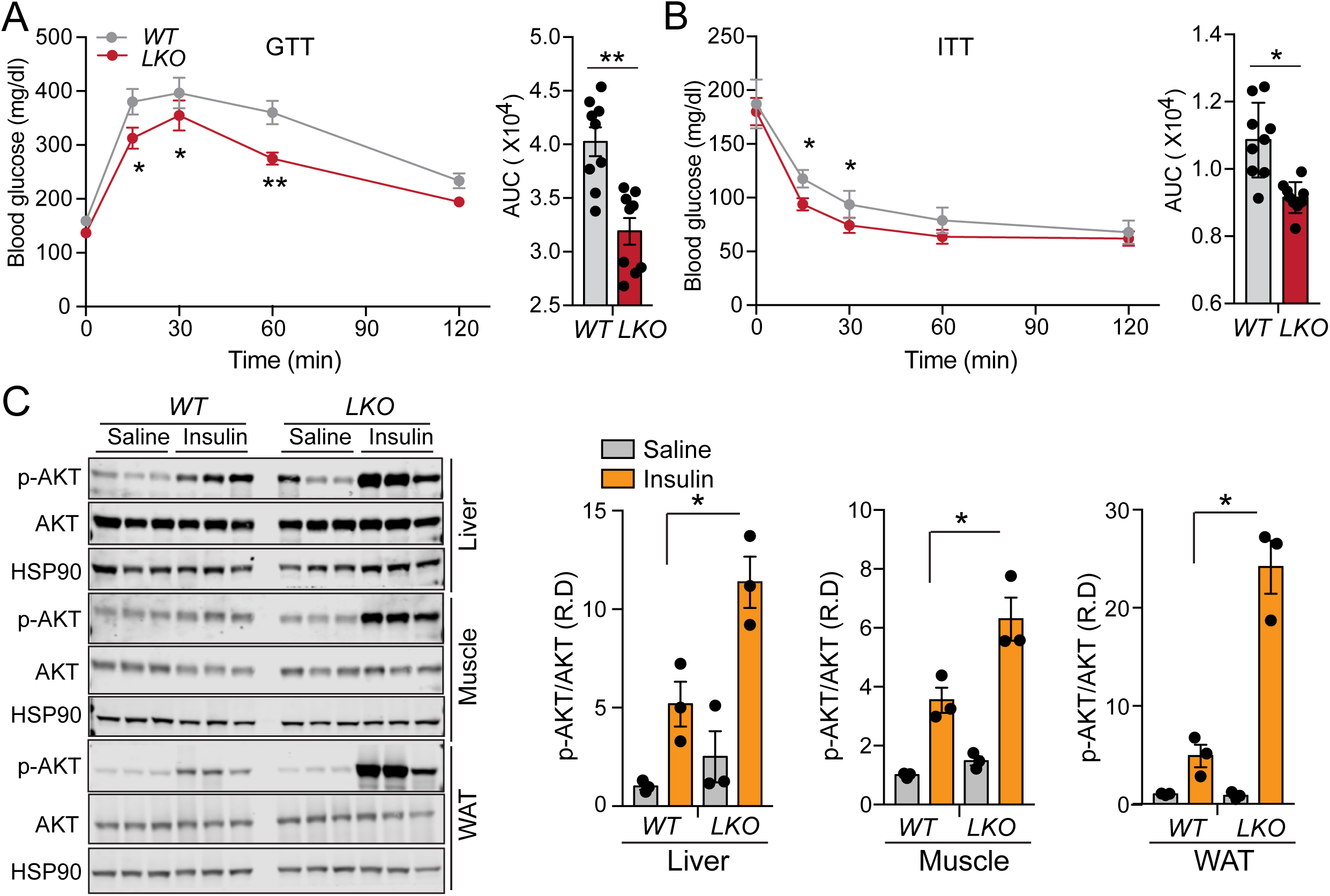
Lack of ANGPTL4 in the liver exhibits improved glucose homeostasis and enhanced insulin sensitivity. (**A**) Intraperitoneal glucose (1 g/kg body weight) tolerance test (GTT) and area under the curve (right panels) of mice fed an HFD for 16 weeks. (**B**) Intraperitoneal insulin (2.5U/kg body weight) tolerance test (ITT) and area under the curve (right panels) of mice fed an HFD for 16 weeks. (**C**) Representative immunoblot images and quantification of Akt (Ser 473 phosphorylation status relative to total AKT (right panels) in the liver, muscle, and adipose tissue, 15 min after an intraperitoneal bolus of insulin (2.5 U/kg) or saline in 16 weeks HFD fed WT vs. *LKO* mice. R.D denotes relative density. All data represent mean ± SEM. *p<0.05 **p<0.01 comparing *LKO* with WT mice using the unpaired t-test.

### Reduced body weight and circulating lipids are associated with attenuated atherosclerosis in hypercholesterolemic mice

Elevated levels of apoB-containing lipoproteins is a major risk factor for the development of atherosclerosis in mice and humans (36, 37). As we have observed that hepatic loss of ANGPTL4 resulted in reduced circulating lipids, we assessed whether accelerated clearance of TRL-remnants reduces atherosclerosis severity in *LKO* mice. To this end, *LKO* and WT mice were injected with the AAV8-PCSK9 adenoviral vector encoding a gain-of-function mutation in PCSK9, to degrade the LDL receptor (LDLR) and induce hyperlipidemia/atherosclerosis. Two weeks following the injection, mice were fed a WD (40% fat Kcal, 1.25% cholesterol) for 16 weeks, and body weight and food intake were monitored weekly. At the end of the study, metabolic parameters and atherosclerotic burden were assessed. Consistent with the phenotype observed in HFD fed mice, we observed that *LKO* mice were resistant to weight gain with no change in food intake upon WD feeding, which was reflected in reduced fat mass as compared to WT (**Figure 5A**; **Figure S3A** and **B**). While circulating HDL-C levels were unaltered, we observed significantly reduced circulating TAGs, TC, and fasting blood glucose in LKO mice after WD feeding (**Figure 5B**; **Figure S3C** and **D**). FPLC analysis of plasma lipoprotein fractions revealed decreased plasma TAG and cholesterol levels in VLDL fractions and cholesterol in IDL/LDL fractions of *LKO* mice (**Figure 5C**).

**Figure 5.**
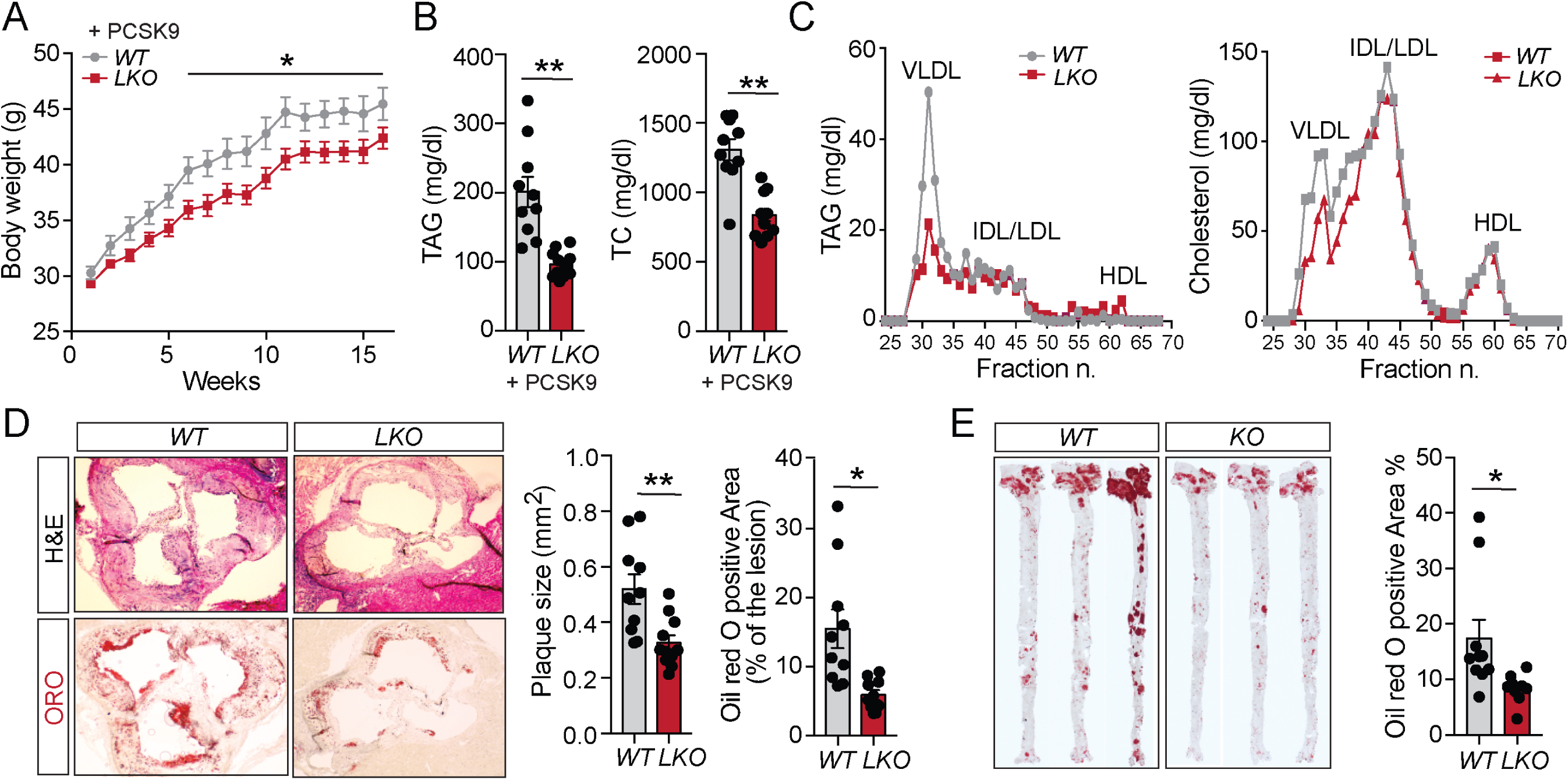
Genetic Ablation of ANGPTL4 in the liver improves obesity and attenuates atherosclerosis. (**A**) Body weight (**B**), Plasma triacylglycerides (TAG), and total cholesterol (TC) from WT and *LKO* mice fed a western-type diet (WD) for 16 weeks. (**C**) Lipoprotein profile analysis of pooled plasma of WT and *LKO* mice (n=5). (**D**) Left: representative histological analysis of a cross-section of the aortic root sinus isolated from WT & *LKO* mice fed a WD for 16 weeks stained with H&E (upper panels) and Oil red O (ORO; lower panels). Right: quantification of plaques size and the percentage area of neutral lipid accumulation calculated from H&E or ORO cross-sections, respectively. (**E**) Representative pictures from the *en face* analysis of aorta isolated from WT and *LKO* mice fed a WD for 16 weeks. All data represent mean ± SEM. *p<0.05, **p<0.01 comparing *LKO* with WT mice using the unpaired t-test.

Similar to what we observed in mice lacking ANGPTL4 in adipose tissue, *LKO* mice also exhibited significantly reduced plaque area in the aortic root as compared to WT littermates (**Figure 5D**). As indicated by Oil-Red-O staining, plaque lipid accumulation was markedly reduced in the *LKO* mice (**Figure 5D**). Similarly, en-face Oil-Red-O staining showed a significantly decreased lipid accumulation in the whole aorta of *LKO* mice (**Figure 5E**). The reduced lipid deposition in the aorta of *LKO* mice was accompanied by a marked reduction in vascular inflammation, as shown by the significant decrease in macrophage accumulation in the lesions (**Figure S3E**). However, we did not observe any significant difference in the circulating blood leukocytes between the groups (**Figure S3F**).

### Liver-specific loss of ANGPTL4 suppresses endogenous lipogenic pathway and promotes oxidative metabolism

To investigate potential mechanisms by which loss of ANGPTL4 in the liver reduces hepatic lipid accumulation despite increased lipid uptake, we analyzed the expression and activity of the main enzymes (ACC, FASN, and HMGCR) involved in *de novo* lipogenesis (DNL)(38, 39). Both the expression and activity of ACC, FASN, and HMGCR were significantly reduced in the liver of *LKO* mice as compared to WT mice fed with CD (**Figure 6A-C**). Next, we assessed whether FAO is altered in the liver as a consequence of depletion of ANGPTL4. We observed markedly increased hepatic FAO in *LKO* mice as compared to WT mice fed CD (**Figure 6D**). We also observed similar results in *LKO* mice fed HFD/WD diet (**Figure S4A-L**).

**Figure 6.**
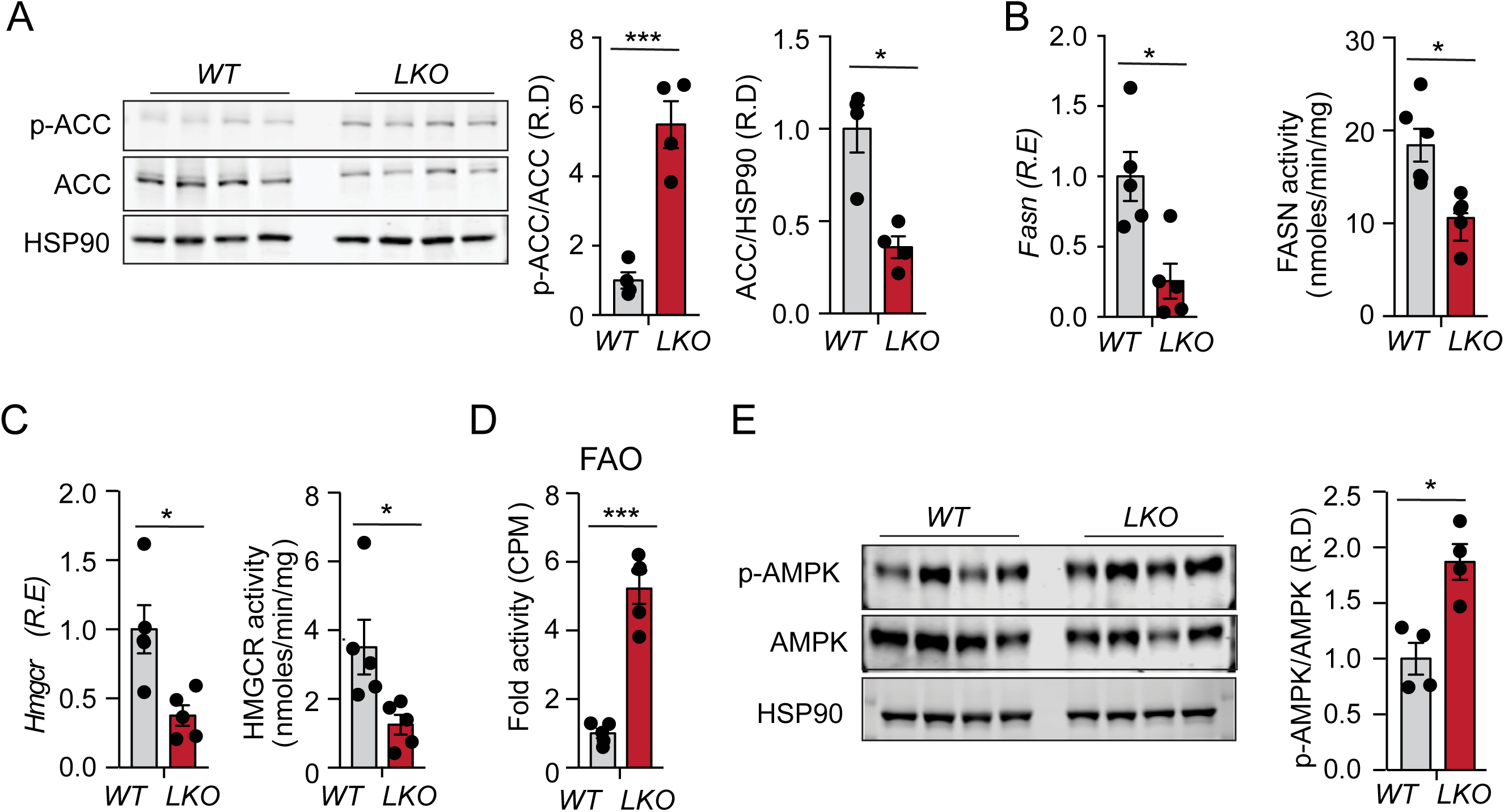
Loss of ANGPTL4 in the liver inhibits DNL pathway and promotes FAO. (**A**) Representative immunoblot images showing p-ACC and ACC levels in the liver isolated from fasted WT and *LKO* mice fed a chow diet (CD) for eight weeks. The right panel shows p-ACC/ACC & ACC/HSP90 ratios from immunoblot images quantification. (**B-D**) Expression and the respective activity of the enzymes FASN and HMGCR and FAO in the liver isolated from fasted mice. (**E**) Immunoblot images of p-AMPK and total AMPK protein in the liver isolated from fasted mice. Densitometric analysis shown in the right panels. R.E denotes relative expression and R.D denotes relative density respectively. All data represent the mean ± SEM. *p<0.05 ***p<0.001 comparing *LKO* with WT mice using the unpaired t-test.

Previously it has been shown that ANGPTL4 might regulate AMPK activity in the hypothalamus (40). Moreover, since AMPK is known to coordinate FA partitioning between oxidation and biosynthesis pathways by increasing FAO capacity and inhibiting DNL, we investigated whether AMPK activity in the liver of *LKO* mice was increased owing to the enhanced lipid uptake. Notably, we observed a significant increase in the phosphorylation of AMPK in the liver of *LKO* mice as compared to WT (**Figure 6E**). To further understand the mechanism of activation of AMPK in the absence of ANGPTL4, we silenced the expression of ANGPTL4 in the human hepatoma HepG2 cell line. Consistent with the *in vivo* findings, shRNA-mediated silencing of ANGPTL4 in HepG2 cells (HepG2-shANGPTL4) (**Figure 7A** and **B**) resulted in enhanced phosphorylation of AMPK and ACC as compared to control (HepG2-shC) (**Figure 7C**, lines one and four). Since absence of ANGPTL4 promotes FAO, and increased FAO is known to promote ROS generation (41), we hypothesized that the enhanced AMPK activation is mediated by ROS. To test this hypothesis, we first analyzed ROS levels in HepG2-shANGPTL4 cells. The results showed that ROS production was significantly increased in HepG2-shANGPTL4 cells as compared to HepG2-shC cells (**Figure 7D**). Most importantly, reducing ROS levels using N-acetyl cysteine (NAC) (**Figure 7D**), attenuated AMPK and ACC phosphorylation (**Figure 7C**, lines three and six) and decreased FAO (**Figure 7E**) in HepG2-shANGPTL4 cells. We further assessed whether AMPK mediates the effects of ROS on regulation of FAO by ANGPTL4 using compound C (CompC), a well-established inhibitor of AMPK. Following treatment with CompC, no differences were observed in AMPK phosphorylation (**Figure 7C**, lines four and five) or FAO (**Figure 7E**) in ANGPTL4 deficient HepG2 cells. These results correlate with the effects of NAC and CompC on the expression of genes regulating FAO (**Figure 7F**) and lipid synthesis (**Figure 7G**). Taken together, these findings demonstrate that absence of ANGPTL4 in hepatic cells increases ROS production leading to AMPK activation, which further enhances FAO and suppresses lipogenesis.

**Figure 7.**
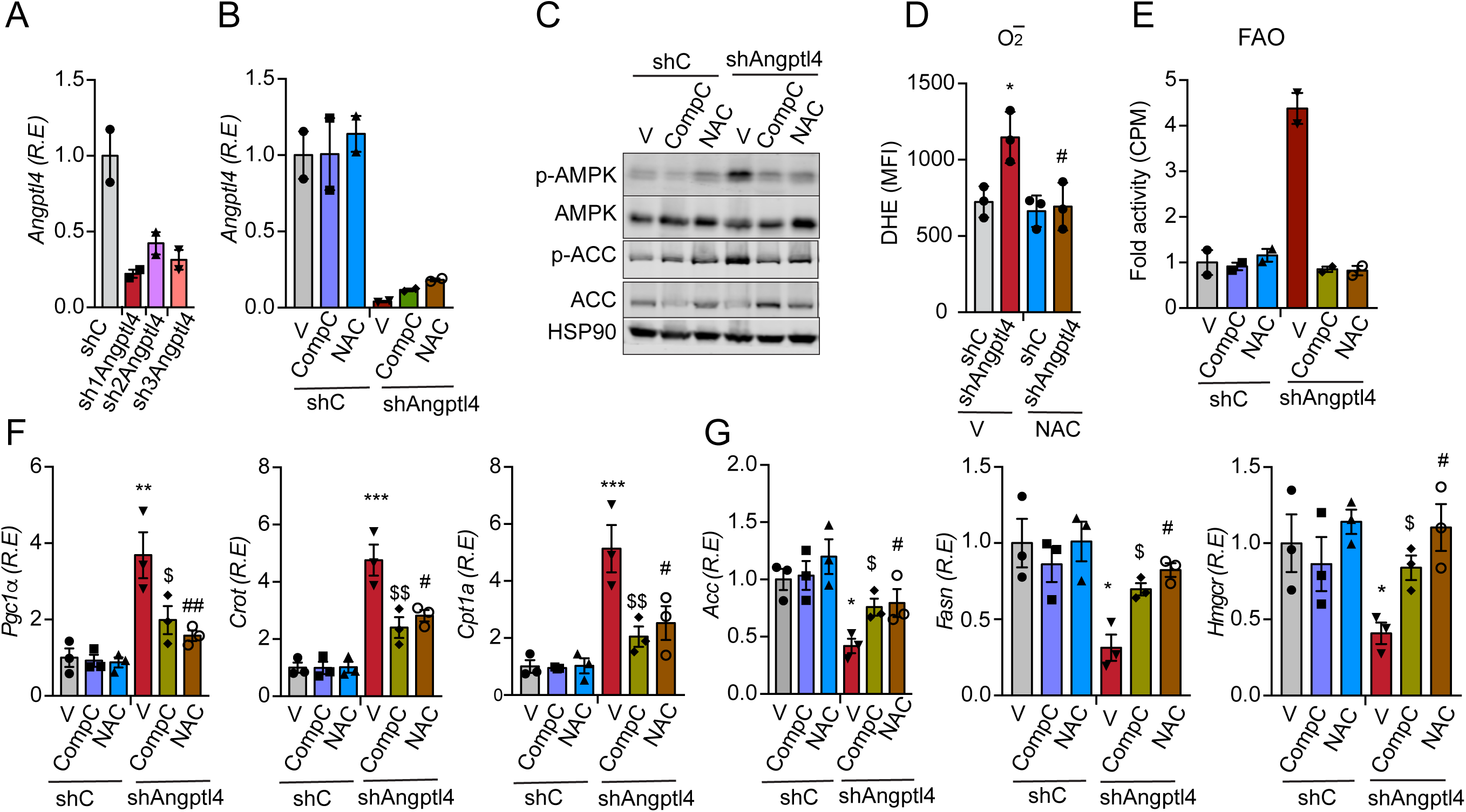
Inhibition of ROS and AMPK in HepG2 cells abrogates the effect of ANGPTL4 in lipid metabolism. (**A**) HepG2 cells were stably transfected with 3 different shRNAs against *Angptl4*. Validation of knockdown efficiency of shRNAs against *Angptl4* in HepG2 cells via qRT-PCR. (**B**) Cells were treated with either NAC (5 mM) or compound C (20 μM) for 24h and evaluation of expression of *Angptl4*. (**C**) Representative immunoblot images showing the levels of p-AMPK, AMPK, p-ACC, and ACC in HepG2-shC and HepG2-shNAGPTL4 cells grown under the similar conditions mentioned in (**B**). (**D**) Relative ROS generation in HepG2-shC and HepG2-shNAGPTL4 cells upon treatment with NAC for 24h. (**E**) FAO in cells under the condition mentioned in (**B**) (**F**) Gene expression of *Pgc-1α, Crot*, and *Cpt1a* in HepG2-shC and HepG2-shNAGPTL4 cells. (**G**) mRNA levels of *Acc, Fasn*, and *Hmgcr* in HepG2-shC and HepG2-shNAGPTL4 cells. R.E denotes relative expression. All data represent mean ± SEM. *,$,#p<0.05, **,$$,##p<0.01, ***p<0.001 as determined by two way ANOVA followed by Bonferroni posttest analysis (* represent comparison between HepG2-shANGPTL4 and HepG2-shC while # and $ represent comparison between HepG2-shANGPTL4 –vehicle ctrl vs HepG2-shANGPTL4-CompC and HepG2-shANGPTL4 –vehicle ctrl vs HepG2-shANGPTL4 -NAC respectively).

### Inhibition of AMPK and ROS production in the liver abrogates the beneficial effect of liver-specific deletion of ANGPTL4

To validate these *in vitro* findings and assess whether ROS mediated AMPK activation is involved in mediating the effects of ANGPTL4 deficiency on DNL and FAO, mice were administered with either CompC or NAC. Consistent with the *in vitro* data, treatment with NAC reduced ROS levels in primary hepatocytes isolated from NAC treated *LKO* mice (**Figure 8A** and **B**). Following treatment with CompC or NAC, we no longer observed any differences in the phosphorylation of AMPK and ACC (**Figure 8C**), FAO (**Figure 8D**), or the expression of genes involved in FAO (*Pgc-1α, Cpt1a, Crot*, and *Hadhb*) between *LKO* and control mice (**Figure 8E**). Similarly, the expression of genes regulating DNL, including *Acc, Fasn*, and *Hmgcr*, were not altered in *LKO* mice treated with CompC as compared to controls (**Figure 8F**), nor was the activity of FASN and HMGCR (**Figure 8G** and **H**). Collectively, both *in vitro* and *in vivo* data demonstrate that inhibition of ROS mediated AMPK activation in the liver reversed the beneficial metabolic phenotype observed in the absence of hepatic ANGPTL4.

**Figure 8.**
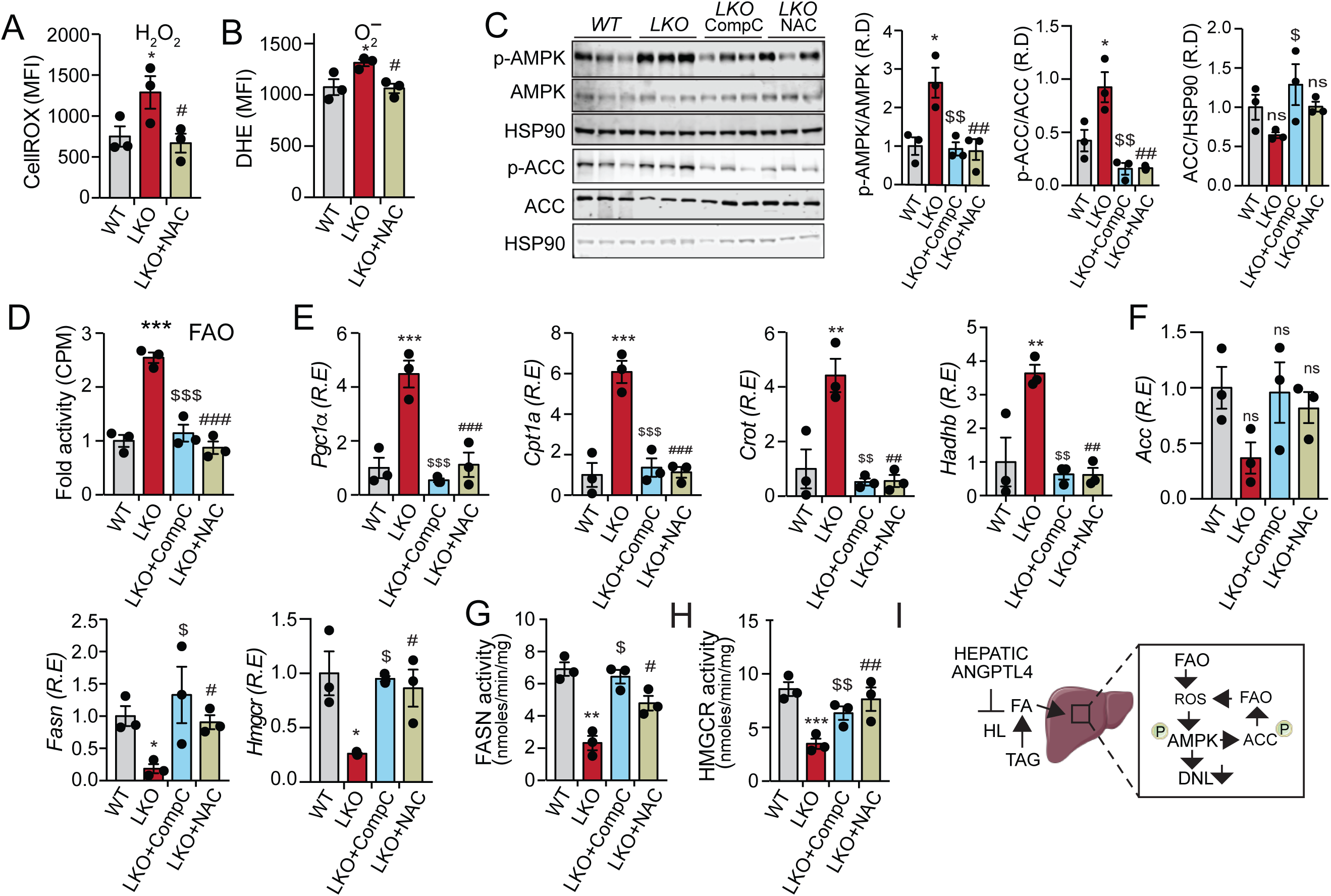
Inhibition of ROS dependent activation of AMPK in *LKO* mice reverses the hepatic lipid metabolism. Eight weeks old *LKO* mice were divided randomly into three groups. Each group of mice was administered with vehicle control (V), compound C (CompC), and NAC, respectively, for the three consecutive days. (**A** and **B**) Determination of cellular ROS (H_2_O_2_ and O_2_^-^) in the hepatocytes isolated from the *LKO* mice with indicated treatment groups. (**C**) Immunoblots showing the levels of p-AMPK, AMPK, p-ACC, and ACC in the liver isolated from fasted *LKO* mice from the indicated treatment groups. The right panel shows image quantification of p-AMPK/AMPK, p-ACC/ACC, and ACC/HSP90 ratios from immunoblot densitometry. (**D**) FAO in the liver of fasted *LKO* mice from indicated treatment groups. (**E**) mRNA expression profile of FAO genes and (**F**) FA biosynthesis genes in the liver of the mice administered with indicated inhibitors, as accessed by qRT-PCR. (**G** and **H**) FASN and HMGCR enzymatic activity in the liver. (**H**) Proposed mechanism for the role of liver-derived ANGPTL4 in hepatic lipid metabolism. R.E denotes relative expression and R.D denotes relative density respectively. All data represent mean ± SEM. *,$,#p<0.05, **,$$,##p<0.01, ***,$$$,###p<0.001 as determined by one way ANOVA followed by Bonferroni posttest analysis (* represent comparison between WT and *LKO* mice while # and $ represent comparison between *LKO*-ctrl vs *LKO*-CompC and *LKO*-ctrl vs *LKO*-NAC respectively).

### GalNac-conjugated ANGPTL4 ASO treatment improves body weight, fasting plasma TAG and insulin resistance in HFD-fed mice

Previous studies in humans have shown that loss of function mutations in the *ANGPTL4* locus are associated with reduced type 2 diabetes and risk of cardiovascular disease (21, 25). However, mice lacking ANGPTL4 and fed a high fat diet enriched in saturated fat develop gut inflammation, characterized by the accumulation of lipid-laden macrophages. Additionally, mice and monkeys treated with neutralizing antibodies against ANGPTL4 develop moderate systemic inflammation. These deleterious effects created a roadblock to the use of ANGPTL4-based therapies. Given the beneficial metabolic effects observed in mice lacking ANGPTL4 in the liver, we evaluated whether a targeted therapy that inhibits ANGPTL4 specifically in the liver might overcome the adverse effects found in the global ANGPTL4 deficient mice and mice treated with ANGPTL4 neutralizing antibodies. To this end, we treated mice with N-acetyl galactosamine conjugated antisense oligonucleotides against ANGPTL4 (GalNac-conjugated ASO). These constructs have a high-affinity for the hepatocyte-specific asialoglycoprotein receptor, therefore its conjugation allows specific inhibition of a target gene in the liver (42). Ten-week old male C57BL6 mice were administered with GalNac-conjugated ANGPTL4 ASOs (ANGPTL4 ASO) or GalNac-control ASOs (Ctrl ASO) via retro-orbital injections, once a week for six weeks under CD fed conditions (**Figure 9A**). After six weeks of treatment, the expression of *Angptl4* mRNA was decreased by ∼ 70% in the liver but was not changed in the epididymal fat (**Figure 9B**), suggesting its ability to inhibit ANGPTL4 explicitly in the liver.

**Figure 9.**
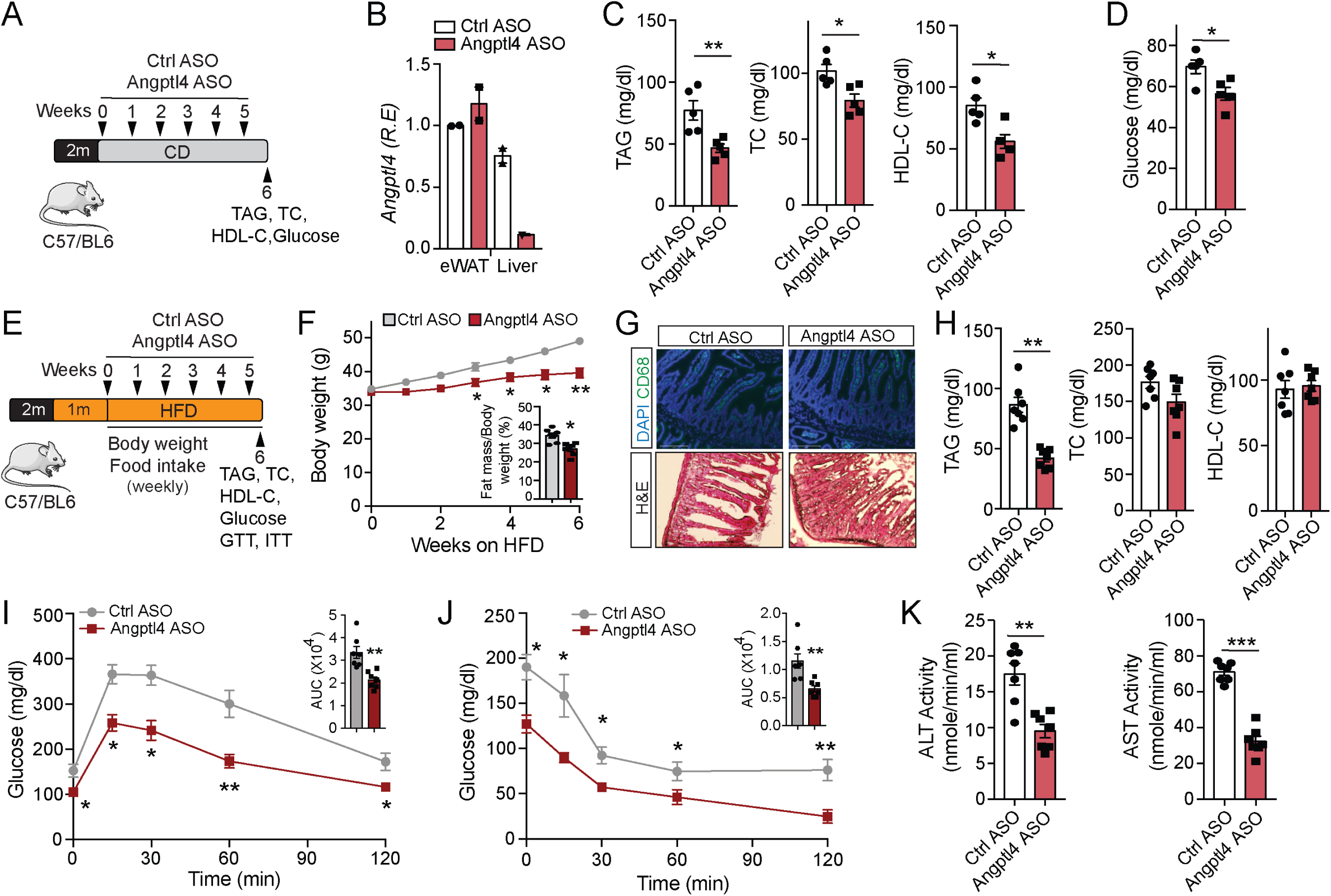
GalNac-conjugated ANGPTL4-ASO treatment improves whole body metabolism in physiological and pathophysiological conditions. (**A**) Schematic presentation of the experimental design of GalNac-conjugated ANGPTL4-ASO (ANGPTL4 ASO) treatment of chow diet (CD) fed mice. (**B**) *Angptl4* expression in eWAT and liver. (**C**) Plasma TAG, TC and HDL-C from ANGPTL4 ASO or Ctrl-ASO treated WT mice. (**D**) Fasting blood glucose (**E**) The strategy for the treatment of GalNac-conjugated ANGPTL4 ASO in fat-induced obese mice. HFD fed mice were treated with ANGPTL4-ASO or Ctrl-ASO for 6 weeks. (**F**) Body weight: the number of weeks on a HFD diet is indicated (The treatment was started at week 5 of HFD feeding). Inset represent the fat mass measured by Echo-MRI. (**G**) Representative images of small intestine cross-sections of HFD fed mice from the 10-week treatment of ANGPTL4 ASO or Ctrl ASO were stained with macrophage marker CD68 and H&E. (**H**) Plasma TAG, TC and HDL-C from ANGPTL4 ASO or Ctrl ASO treated HFD fed WT mice. (**I** and **J**) Intraperitoneal glucose tolerance test (GTT) and Intraperitoneal insulin tolerance test (ITT) in ANGPTL4 ASO or Ctrl ASO injected in mice fed a HFD. Inset represents the AUC. (**K**) The activity of plasma ALT and AST after treatment of ANGPTL4 ASO or Ctrl ASO in HFD-induced obese mice. All data represent mean ± SEM. *p<0.05 **p<0.01 ***p<0.001 comparing ANGPTL4 ASO with Ctrl ASO treated mice using the unpaired t-test.

Administration of the ANGPTL4 ASO resulted in decreased circulating levels of TAG, TC, HDL-C and fasting glucose, as well as a reduction in TAG within VLDL and IDL/LDL fractions and cholesterol within HDL-C fractions (**Figure 9C** and **D**; **Figure S5B** and **C**). However, body weight was unaltered between both groups of mice (**Figure S5A**).

To further determine whether therapeutic inhibition of ANGPTL4 in the liver attenuates HFD-induced obesity and insulin resistance, mice were fed a HFD for 4 weeks followed by administration of ANGPTL4 ASO for six weeks (**Figure 9E**). We monitored body weight and food intake weekly, and after the six week of treatment lipid parameters and glucose homeostasis were analyzed. Notably, ANGPTL4 ASO treatment significantly decreased body weight and body-fat percentage in HFD fed mice (**Figure 9F**). As global ANGPTL4 KO mice are shown to develop severe systemic metabolic complications after ∼10 weeks on HFD diet (26), we analyzed gut inflammation and circulating leukocytes after 10 weeks of treatment of ANGPTL4 ASO. Treatment with ANGPTL4 ASO had no considerable effect on food intake, gut inflammation, or circulating leukocytes in mice after 10 weeks of treatment (**Figure 9G**; **Figure S5D** and **E**), suggesting that the reduction in body weight of ANGPTL4 ASO treated mice was independent of food intake or gut inflammation.

Furthermore, ANGPTL4 ASO treatment resulted in decreased fasting plasma TAGs in mice fed a HFD (**Figure 9H**). Plasma lipoprotein fraction analysis showed a reduction TAGs in both the VLDL and IDL/LDL fractions after GalNac-Angptl4 ASO treatment (**Figure S5F**). Plasma TC and HDL-C remain unchanged in both the treatment groups (**Figure 9H**; **Figure S5G**). To evaluate the effect of ANGPTL4 silencing on glucose homeostasis, we performed GTT and ITT in mice fed a HFD for 6 weeks and treated with ANGPTL4 ASO. We observed a significant improvement in glucose tolerance and insulin sensitivity in mice administered with ANGPTL4 ASO (**Figure 9I** and **J**). Moreover, we found a significant reduction in the liver weight and hepatic TAG levels in ANGPTL4 ASO treated mice as compared to Ctrl-ASO treated mice (Figure S5H). Importantly, ANGPTL4 ASO treatment appeared to protect against HFD induced liver damage, as plasma levels of liver enzymes alanine aminotransferase (ALT) and aspartate aminotransferase (AST) were reduced significantly in ANGPTL4 ASO treated obese mice as compared to obese mice treated with Ctrl ASO (**Figure 9K**). Together, these data indicate that silencing of ANGPTL4 in the liver using GalNac conjugated ASOs recapitulates the beneficial effects of LKO mice in both physiological as well as pathological conditions, notably without developing any of the systemic metabolic abnormalities observed in ANGPTL4 global knockout or ANGPTL4 antibody treated mice. More importantly, GalNac conjugated ANGPTL4 ASOs were able to restore metabolic function in animals that were already obese, suggesting that this may provide a viable treatment option for obesity related metabolic disorders, including T2D and cardiovascular disease.

## DISCUSSION

Elevated circulating levels of ANGPTL4 and TAGs are key etiologic components of metabolic disorders like obesity, T2D, and CAD, which are associated with increased adiposity, fatty liver, and insulin resistance. Human studies indicate a positive relationship between ANGPTL4 function and body mass index (BMI), fat mass, glucose intolerance, insulin resistance, and circulating lipids (25). These observations strongly suggest that ANGPTL4 could be a potential target in metabolic diseases. In this line, many efforts have been made to study the role of ANGPTL4 in metabolic diseases. However, the exact functions are not yet fully understood because of the lack of knowledge of the cell/tissue-specific roles of ANGPTL4. This is especially relevant because severe systemic metabolic complications (i.e., gut inflammation) confound the researcher’s ability to investigate the beneficial functions of ANGPTL4 in whole-body KO mice upon HFD/WD feeding (26, 31). Since ANGPTL4 is highly expressed in the liver and AT of both humans and mice, we generated ANGPTL4 knockout mouse models specific to these issues. We have previously reported that adipose-specific ANGPTL4 knockout (Ad-KO) mice exhibit improved plasma lipid profiles and insulin sensitivity, but no difference in body weight between Ad-KO and WT mice during prolonged HFD feeding (32). As the overall impact on systemic metabolism was not as profound as expected, we hypothesized that ANGPTL4 derived from another vital metabolic organ involved in lipid homeostasis might be responsible for the effects on whole-body lipid and glucose metabolism observed in human studies. Interestingly, the ablation of hepatic ANGPTL4 resulted in decreased body weight, plasma lipids (TAGs and TC), hepatic steatosis, and improved glucose tolerance and insulin sensitivity after 16-weeks of HFD feeding. The effects observed in *LKO* mice were more pronounced than those found in Ad-KO mice, indicating that the deficiency of ANGPTL4 in the liver has major consequences on systemic lipid and glucose metabolism.

The improved circulating lipids in liver-specific ANGPTL4 KO mice was accompanied by accelerated TRL-remnant catabolism. We observed increased plasma TAG clearance and enhanced HL activity in *LKO* mice. However, conflicting reports exist on the regulation of HL activity by ANGPTL4. Koster *et al*., using global ANGPTL4 KO mice and liver-specific overexpression of ANGPTL4, and Gutsell e*t al*., using an *in vitro* system, have shown that ANGPTL4 does not affect HL activity (28, 43). In contrast, Lichtenstein *et al*. observed that adenovirus-mediated overexpression of ANGPTL4 inhibits post-heparin HL activity (12). The cause for this discrepancy may be related to the differences in the mouse models used (magnitude of ANGPTL4 expression) and sensitivity of the assay used for *in vitro* and systemic plasma HL activity. Increased activity of LPL likely also contributes to the reduced circulating TAGs due to decreased levels of VLDL, consistent with existing data from numerous *in vivo* and *in vitro* studies. However, strikingly, our data suggest that much of the systemic changes in lipid homeostasis in the absence of hepatic ANGPTL4 may be mediated by increased HL activity.

It has been previously reported that HL, in addition to acting as a lipolytic enzyme, also facilitates the uptake of chylomicrons and VLDL remnants by hepatocyte cell surface receptors, thus lowering circulating lipids (44, 45). Comparison of FPLC plasma lipoprotein cholesterol profiles (IDL/LDL-fraction) in atherogenic mice further indicates that the protective impact of hepatic depletion of ANGPTL4 involves the contribution of elevated HL activity. Increased HL activity could facilitate the uptake of TRL-remnants in hepatocytes to prevent the accumulation of atherogenic lipoproteins in atherosclerotic plaques. We also noticed that the depletion of hepatic ANGPTL4 leads to AMPK-mediated suppression of HMGCR expression and activity. Therefore, we believe that the suppressed activity of HMGCR in *LKO* mice might contribute to reduced atherogenic lipoproteins and cholesterol in the plasma. These data suggest that hepatic ANGPTL4 may have a unique role in regulating both HL and HMGCR activity during the progression of atherosclerosis.

In contrast to our prior work in Ad-KO mice, we found that loss of hepatic ANGPTL4 attenuates body weight gain upon HFD feeding, mainly by reducing visceral fat mass and liver weight, without affecting food intake. The reduction in fat mass and size of adipocytes in *LKO* mice could be due to the accelerated hydrolysis of TRL remnants and increased FA uptake in the liver, decreasing circulating TAGs and thereby reducing lipid storage in AT. The role of ANGPTL4 in the regulation of body weight in mice has been controversial under both physiological as well as pathophysiological conditions (13, 21, 28, 32). The different outcomes from these studies may be due to the inactivation of different exons of the N-terminus of ANGPTL4 in the different knockout models, different types of diet, and the magnitude of overexpression. Structure-function analysis of ANGPTL4 from various knockout models through protein crystallography or Cryo-EM may eventually justify these discordant findings.

Despite increased lipid uptake in the liver, we found a marked reduction in the intrahepatic lipid accumulation. A concomitant increase in the rate of FAO in the liver suggests excess fat taken up by the liver is dissipated through enhanced β-oxidation via activation of AMPK. AMPK is known to increase FAO and suppresses DNL by inhibiting ACC. This may be one of the mechanisms by which hepatic deletion of ANGPTL4 improves metabolic function following HFD-induced obesity. This hypothesis is supported by the recent studies on the overexpression of AMPK in the mouse liver (46, 47). Data from these studies indicate that activation of AMPK in the liver decreases lipid deposition not only in the liver, but also in the AT. We also provide evidence that lack of ANGPTL4 in the liver improves whole-body glucose homeostasis and insulin action even upon prolonged HFD feeding, which is likely due to the marked reduction in lipid accumulation in the liver and adipose tissue.

In the present study, we demonstrate a novel mechanism by which ANGPTL4 regulates lipid homeostasis via a mechanism involving HL/ROS/AMPK axis. Increased lipase-mediated lipid uptake causes a compensatory increase in FAO (41), which leads to the ROS induced sustained activation of AMPK. Our data indicate that suppression of ROS-mediated AMPK activation is sufficient to reverse the metabolic phenotype observed in *LKO* mice. ROS at a physiological level play a key role the cellular signaling as well as regulating metabolism (48). Our data suggest that reducing ROS with scavenger NAC not only suppressed activation of AMPK but also reversed changes in hepatic lipid metabolism in *LKO* mice. These observations suggest that ROS might link the inverse relationship between AMPK and ANGPTL4.

In the recent years, studies have shown that ANGPTL4 act as potent metabolic regulator and is strongly associated with various metabolic disorders. Attempts to inhibit ANGPTL4 via using the neutralizing antibody in humanized mice and non-human primates were undermined by severe systemic metabolic abnormalities (22). Our data from *LKO* mice support the hypothesis that liver-specific suppression of ANGPTL4 might have therapeutic potential. Therefore, we evaluated whether therapeutic inhibition of ANGPTL4 in the liver recapitulates the metabolic phenotype observed in *LKO* mice. Interestingly, ANGPTL4 ASO treated mice were ameliorated from HFD-feeding-induced obesity and had improved circulating TAGs, glucose tolerance, and insulin sensitivity. Conclusively, this study reveals that ANGPTL4-ASO mediated inhibition of hepatic ANGPTL4 consistently replicates most of the aspects of the obesity-resistant phenotype observed in *LKO* mice. Most importantly, mice treated with ANGPTL4 ASO did not exhibit any of the systemic metabolic abnormalities observed in previous studies. Thus, these data provide a strong rationale for following the development of liver-specific ANGPTL4 therapeutic agents for the treatment of metabolic diseases.

In conclusion, we show that the absence of ANGPTL4 in the liver prevented obesity-linked diabetes and ameliorated pathological manifestations resulting from chronic HFD/WD feeding in mice via a mechanism involving HL/ROS/AMPK (Figure 8I). This includes reductions in hyperlipidemia, glucose intolerance, insulin resistance, hepatic steatosis, and progression of atherosclerosis. This study not only signifies the contribution of hepatic ANGPTL4 in glucose and lipid homeostasis but also provides new insight into ANGPTL4 and NAFLD-associated diabetes and atherosclerosis susceptibility. Remarkably, our data demonstrate that liver-specific targeting of ANGPTL4 could provide a viable therapeutic approach for the treatment of obesity-induced diabetes and atherosclerosis. As previous work has shown that global loss or inhibition of ANGPTL4 can have profoundly adverse effects on the health and function of the gut, this could provide a substantially less risky approach for the treatment of common chronic metabolic conditions.

### Limitations

While the findings of this study, along with our prior work has considerably improved our understanding of how ANGPTL4 in vital metabolic organs impacts the ability to maintain metabolic homeostasis, there are still important questions and caveats that will need to be addressed in the future. Most importantly, ANGPTL4 is a secreted protein, and our current work has been unable to determine how the deficiency in the liver or AT might affect circulating levels of ANGPTL4 and what impact this may have on lipid metabolism in other tissues. To further explore this, a specific and reliable antibody against mouse ANGPTL4 needs to be generated to measure serum ANGPTL4 levels. Also, we have used a pharmacological inhibitor of AMPK, CompC, which might have off-target effects. Therefore, a genetic approach to silence AMPK in the liver would be more appropriate approach.

## MATERIALS AND METHODS

### Animal studies

Generation of liver-specific ANGPTL4-deficient mice. Mice bearing a loxP-flanked ANGPTL4 allele (*ANGPTL4*^*loxP/loxP*^ mice) were generated as described previously (32). Liver-specific ANGPTL4–deficient mice (Albumin-Cre; *ANGPTL4*^*loxP/loxP*^, *LKO*) were generated by breeding albumin-Cre; *ANGPTL4*^*loxP/*+^ mice with *ANGPTL4*^*loxP/*+^ mice. All mouse strains were in the BL6 genetic background. *LKO* mice were verified for Angptl4 KO in the liver by PCR using Cre primers and primers flanking the 5′ homology arm of the ANGPTL4 gene and LoxP sites from the tail-extracted DNA. All experimental mice were housed in a barrier animal facility with a constant temperature and humidity in a 12-hour dark/light cycle while water and food were provided *ad labium*. All mice (n=3-5 per cage) were fed with a standard chow diet (CD) for 8 weeks after that switched to an HFD (60% calories from fat; Research Diets D12492) for 1–16 weeks. For studying atherosclerosis, mice (8-week-old, n=3-5 per cage) were administered with a single retro-orbital Injection of recombinant adeno-associated virus (AAV, containing 1.0 × 10^11^ genome copies) encoding PCSK9 (AAV8.ApoEHCR-hAAT.D377Y-mPCK9.bGH) to induces hyperlipidemia. Two weeks’ post-injection, atherosclerosis was induced by feeding the mice with high cholesterol, western diet (WD) containing 1.25% cholesterol (D12108, Research Diets). Body weight and food intake were measured every week of HFD and WD fed mice.

For liver-specific *Angptl4* antisense oligonucleotide (ASO) studies, mice, at ten weeks of age, were injected with GalNac-control ASO (Ctrl ASO) or GalNac conjugate *Angptl4* ASO (Angptl4 ASO) (the combination of two ASOs) through retro-orbital rout at a dose of 25mg/kg/week for six weeks. ASO sequences used in this study were as follows: Angptl4 ASO (1) 5’-AGCTGTAGCAGCCCGT-3’ (2) 5’-ATATGACTGAGTCCGC-3’ and control ASO 5’-GGCCAATACGCCGTCA-3’. ASOs were dissolved in PBS for the mice experiments. During the GalNac conjugated *Angptl4* ASO treatment, mice were fed with CD. To assess the therapeutic effect of GalN-ASO in obese mice, mice were fed a HFD for 4 weeks and were treated with GalNac-cntr ASO or GalNac-Angptl4 ASO (25mg/kg) by retro-orbital injections weekly for the 10 weeks.

### Glucose and insulin tolerance test

Glucose tolerance tests (GTT) and Insulin tolerance test (ITT) were performed as described previously (32). Briefly, HFD fed *WT* and *LKO* mice fasted for 16 h followed by intraperitoneal injection of glucose (1 g/kg). Glucose levels were determined in the blood collected from the tail vein at indicated time points using a Contour Ultra blood glucometer. Similarly, for determining ITT, 6 h fasted mice were injected with insulin (2.5 U/kg) intraperitoneally, and glucose levels in the blood were analyzed immediately before and after 15, 30, 60, and 120 minutes’ post insulin injection.

### Analysis of insulin signaling *in vivo*

Following 6 h fast, mice were injected (IP) with 2.5 U/kg of regular human Insulin (Novolin, Novo Nordisk). Fifteen minutes post the insulin injection, mice were euthanized according to the approved procedure, and tissues were removed and flash-frozen in liquid nitrogen. The insulin-signaling molecule was analyzed via immunoblotting of p-AKT, as previously described (32).

### Lipoprotein profile and lipid measurements

Blood was collected by tail vein from the overnight-fasted (12-16h) mice, and plasma was separated by centrifugation at 10000 rpm at 4°C for 10 minutes. HDL-C was separated by precipitation of non–HDL-C, and both HDL-C fractions. Total plasma TAGs and cholesterol were enzymatically analyzed using commercially available kits (Wako Pure Chemicals). Distribution of lipids in the plasma lipoprotein fractions was analyzed by FPLC gel filtration with 2 Superose 6 HR 10/30 columns (Pharmacia Biotech).

### Fat tolerance test

A fat tolerance test was performed according to the protocol described previously (49). Briefly, mice fasted for 4 h followed by oral gavage of 10 μl olive oil/gram of body weight. Blood samples were withdrawn from the tail vein post 0, 1, 2, and 4 h administration of olive oil.

### Tissue lipid uptake

Lipid uptake in tissue (s) was performed using a ^3^H labeled triolein, as described previously (49).

### LPL and HL Activity Assay

Post-heparin plasma was collected as previously described (25). Enzymatic activity of hepatic lipase and post-heparin lipoprotein lipase was determined by using ^3^H-labeled triolein according to the protocol described previously (32, 50).

### Hepatic VLDL-TAG secretion

For measuring hepatic VLDL-TAG production rate, mice were fasted overnight (14-16 hours), followed by i.p. Injection with 1 g/kg of body weight poloxamer 407 (Sigma-Aldrich) in PBS. Blood was collected immediately before injection and at 1, 2, 3, and 4 h post-injection as described earlier (32). TAG level was determined using commercially available kits.

### Intestinal lipid absorption

Gut lipid absorption was measured as described earlier (49)(Ref). Briefly, 6 h-fasted mice were injected with 1 g/kg poloxamer 407. Mice were gavaged with emulsion mixture containing 2 μl [^3^H]-triolein, and of 100 μl of mouse intralipid 20% emulsion oil, 1 h post poloxamer injection. Blood was collected at different time points, as indicated. [^3^H]-radioactivity was determined in the plasma.

### Fatty acid oxidation

Ex vivo fatty acid (FA) oxidation was analyzed using [^14^C] palmitate, as described earlier (32).

### FA synthase (FASN) activity assay

FASN activity was determined in the liver, as described previously with some modifications (51). Briefly, liver from *WT* and *LKO* mice were homogenized in tissue homogenization buffer (0.1 M Tris, 0.1 M KCl, 350 mM EDTA, and 1 M sucrose; pH 7.5) containing protease inhibitor cocktail (Roche). The supernatant was collected by centrifuging liver homogenates at 9,400 g for 10 minutes at 4°C. For determining FASN activity, Liver homogenate was added to NADPH activity buffer (0.1 M potassium phosphate buffer, pH 7.5 containing 1 mM DTT, 25 μM acetyl-CoA, and 150 μM NADPH). Malonyl-CoA (50 μM) was added to the assay buffer to initiate the reaction. An increase in the absorbance was followed at 340 nm for 30 min at an interval of 1 min using a spectrophotometer set in the kinetic mode under constant temperature (37°C). FASN activity is represented as n moles NADPH consumed per minute per mg protein.

### Histology, immunohistochemistry, and morphometric analyses

Hearts of mouse were perfused with PBS before the isolation of organs for histology and immunohistochemistry analysis and were incubated in 4% paraformaldehyde for 4 hours. Following the incubation in paraformaldehyde, tissues were three times washed with PBS and incubated with PBS for 30-60 minutes and then kept in 30% sucrose for 12-16 hours at 4°C. Subsequently, hearts were embedded in OCT and frozen. Serial sections of the heart were cut at 6-μm thickness through a cryostat. Every fourth slide from the serial sections was stained with hematoxylin and eosin (H&E), and each consecutive slide was stained with Oil Red O for quantification of the area of the atherosclerotic lesions. Aortic lesion size of each animal was obtained by averaging the lesion areas in at least 9 sections from the same mouse. For the analysis of inflammation in atheroprone areas, 6-μm-thick sections were prepared from the aortic root in the heart from mice that were kept on a WD for 16 weeks. Inflammation in atherosclerotic plaques was assessed by staining with antibodies against CD68 (1:200; Serotec; #MCA1957).

### En face Oil Red O staining

Oil Red O staining was performed as described previously. Briefly, aortas opened up longitudinally were rinsed with 78% methanol, followed by staining with 0.16% Oil Red O solution for 50 minutes, and then destained in 78% methanol for 5 minutes. The lesion area was quantified as a percentage of the Oil Red O–stained area in the total aorta area.

### Liver histology and lipid measurement

The liver samples were either fixed in 4% paraformaldehyde for H&E staining or snap-frozen for oil red O staining. Fixed liver specimens were processed to paraffin blocks, sectioned (5 μm), followed by staining with H&E. For Oil Red O staining, frozen liver samples were embedded in OCT and then sectioned (10 μm) and stained with oil red O to visualize neutral lipids as described previously (21). For determining total TAGs content in the liver, TAGs were extracted using a solvent chloroform/methanol (2:1). TAG level in the liver was determined by using a commercially available assay kit (Sekisui) according to the manufacturer’s instructions.

### ALT and AST measurements

ALT and AST activity was determined in serum with the commercially assay kits (Sigma-Aldrich, MAK052; and MAK055) following manufacturer’s recommendations.

### Circulating leukocyte analysis

Blood was collected by retro-orbital puncture in heparinized microhematocrit capillary tubes. Total numbers of circulating blood leukocytes was measured using a HEMAVET system. For further characterization of leukocyte, FACs analysis was performed. Briefly, erythrocytes were lysed with ACK lysis buffer (155 mM ammonium chloride, 10 mM potassium bicarbonate, and 0.01 mM EDTA, pH 7.4) and leukocytes blocked with 2 μg ml^-1^ of FcgRII/III, followed by staining with a cocktail of antibodies. Monocytes were identified as CD115hi and subsets as Ly6-C^hi^ and Ly6-C^lo^; neutrophils were identified as CD11b^hi^Ly6G^hi^; B cells were identified as CD19^hi^B220^hi^; T cells were identified as CD4^hi^ or CD8^hi^. The following antibodies were used for all the analysis of leucocytes (all from BioLegend): FITC-Ly6-C (HK1.4), PE-CD115 (AFS98), APC-Ly6-G (1A8), PB-CD11b (M1/70), APC-CD19 (6D5), PE/Cy7-B220 (RA3-6B2), APC/Cy7-CD4 (RM4-5) and BV421-CD8a (53-6.7). All antibodies were used at 1:300 dilutions.

### Cell lines and culture conditions

Human liver cell line HepG2 was obtained from American Type Culture Collection (ATCC, USA) and grown in DMEM containing 2 mM/L glutamine supplemented with 10% FBS, penicillin/streptomycin (Life Technologies, USA). HEK293T were maintained in DMEM high glucose supplemented with 10% FBS and 1% penicillin/streptomycin (Life Technologies, USA). Cells were routinely evaluated for mycoplasma contamination.

### Transfection with Plasmids and siRNA

Stable knockdown for Angptl4 was obtained by lentiviral mediated transduction of specific shRNAs against Angptl4 in HepG2 cells and selected in puromycin according to the protocol described earlier (52). A transient knockdown experiment was performed by transfecting siRNAs against Angptl4 and Ampk (sets of 4 SMART POOL siRNAs, Dharmacon, USA) in HepG2 cells using RNAiMAX (Life Technologies, USA) according to the manufacturer’s instructions. Knockdown efficiency was evaluated by qRT-PCR for *Angptl4* and Immunoblots analysis for AMPK and pAMPK.

### Isolation of primary hepatocytes

Primary hepatocytes were isolated from *WT* and *LKO* mice treated with indicated concentrations of inhibitors such as CompC and NAC, according to the protocol described earlier (53).

### Western blot analysis

The liver homogenate was prepared as described previously using the Bullet Blender Homogenizer (31). Both tissues and cells lysates were prepared by lysing in ice-cold buffer containing 50 mM Tris–HCl, pH 7.5, 0.1% SDS, 0.1% deoxycholic acid, 0.1 mM EDTA, 0.1 mM EGTA, 1% NP-40, 5.3 mM NaF, 1.5 mM Na4P2O7, 1 mM orthovanadate, 1 mg/ml protease inhibitor cocktail (Roche), and 0.25 mg/ml AEBSF (Roche). Lysates were further sonicated and rotated at 4°C for 1 hour, followed by centrifugation at 12,000 g for 10 minutes. Lysate protein concentration was determined by the BCA method using a commercially available kit (BIORAD). An equal amount of proteins was resuspended in SDS sample buffer before separation by SDS-PAGE. Proteins were transferred onto a PVDF membranes, and the membranes were probed with the following antibodies: anti-pAMPK, AMPK, anti-pACC, and ACC (AMPK ACC antibody sampler kit, Cell Signaling Technology #9957; dil-1:1,000); AKT, p-AKT, (Cell Signaling Technology #4691 (Akt) #4060T (pAKT S473), dil-1:1,000), and anti-HSP90 (BD Biosciences #610419; dil-1:1,000). Protein bands were visualized using the Odyssey Infrared Imaging System (LI-COR Biotechnology), and densitometry was performed using ImageJ software.

### RNA isolation and quantitative real-time PCR

Total RNA from tissue was isolated using TRIzol reagent (Invitrogen) according to the manufacturer’s protocol. For mRNA expression analysis, cDNA was synthesized using iScript RT Supermix (Bio-Rad), following the manufacturer’s protocol. Quantitative real-time PCR (qRT-PCR) analysis was performed in duplicate using SsoFast EvaGreen Supermix (Bio-Rad) on an iCycler Real-Time Detection System (Eppendorf). The mRNA levels were normalized to 18S.

### NAC and Compound C treatment in mice

To inhibit activation of AMPK and ROS generation *in vivo*, mice were intraperitoneally and orally administered with vehicle control (DMSO), compound C (20 mg/kg) (Tocris) or NAC (1 g/kg in 0.9% saline) (Sigma) respectively for 3 consecutive days. At the end of the experiment, mice were euthanized according to the approved protocol, and liver samples were processed for further studies.

### Measurement of ROS generation

Cellular ROS species H_2_O_2_ and O_2_^-^ was determined in primary hepatocytes isolated from *WT* and *LKO* mice as well as in HepG2 cells using fluorescence dye DCFDA and DHE (Thermo Fisher), according to the manufacturer’s instructions. For measuring mitochondrial ROS, Mitotracker Green (Thermo-Fisher) was used. Briefly, Isolated hepatocytes and cell lines were incubated with 5 μM of DCFDA or DHE or Mitotracker dyes for 30 min at 37°C. Cells were washed twice with PBS, and fluorescence was acquired using a Flow cytometry (FACS Area, BD Bioscience).

### HMGCR Activity Assay

The HMG-CoA reductase activity assay was determined according to the protocol described previously (54), with slight modifications. In Brief, a microsomal fraction from cell lysates and the liver homogenate was obtained via ultracentrifugation (100000 x g for 60 min). The reaction buffer (0.16 M potassium phosphate, 0.2 M KCl, 0.004 M EDTA, and 0.01 M dithiothreitol) containing 100 µM NADPH and microsomal protein (200 µg/mL) was prewarmed at 37 °C for 10 min before the reaction. The reaction was initiated by adding 50 µM substrate (HMG-CoA) to the reaction buffer. The decrease in the absorbance at 340 nm was followed for 30 min with an interval of 1 min. At the same time, a similar assay was performed in the presence of simvastatin, an inhibitor of HMGCR activity. The specific activity of HMGCR was calculated by subtracting total enzyme activity and simvastatin-resistant activity (with simvastatin), and represented as nmol/min/mg of total microsomal proteins.

### Statistics

The mouse sample size for each study was based on literature documentation of similar well-characterized experiments. The number of mice used in each study is listed in the figure legends. In vitro experiments were regularly repeated at least three times unless otherwise noted. No inclusion or exclusion criteria were used, and studies were not blinded to investigators or formally randomized. Data are expressed as average ± SEM. Statistical differences were measured using an unpaired 2-sided Student’s t-test, or 1-way or 2-way ANOVA with Bonferroni’s correction for multiple comparisons. Normality was tested using the Kolmogorov-Smirnov test. A nonparametric test (Mann-Whitney) was used when the data did not pass the normality test. A value of P ≤ 0.05 was considered statistically significant. Data analysis was performed using GraphPad Prism software version 7.

## AUTHOR CONTRIBUTIONS

AKS, BC, YS and CF-H conceived and designed the study and wrote the manuscript. AKS, BC, ACD, XZ, BA, JS, KC and NR performed experiments and analyzed data. NLP and KC analyzed data and edited the manuscript.

## ACKNOWLEDGEMENTS

This work was supported by grants from the National Institutes of Health (R35HL135820 to CF-H; R01HL105945 and R01HL135012 to YS, 5F32DK10348902 to NP, and the American Heart Association (16EIA27550005 to CF-H and 17SDG33110002 to NR), the American Diabetes Association (1-16-PMF-002 to AC-D), and the Foundation Leducq Transatlantic Network of Excellence in Cardiovascular Research MIRVAD (to CF-H).

