## Supplemental Figures for "Liver-specific suppression of ANGPTL4 improves obesity-associated diabetes and mitigates atherosclerosis in mice"

#### SUPPLEMENTAL FIGURE LEGENDS

**Figure S1. Lack of ANGPTL4 in the liver does not influence ANGPTL3 and ANGPTL8 expression in the liver.** Expression of ANGPTL3 and ANGPTL8 in the liver isolated from overnight fasted WT and *LKO* mice as accessed by qRT-PCR. R.E represents relative expression.

**Figure S2. The absence of ANGPTL4 in the liver does not influence food intake but reduces inflammatory marker under HFD fed conditions.** (**A**) Representative image of WT and *LKO* mice fed a HFD for 16 weeks. (**B**) Food intake/day in WT and *LKO* mice fed an HFD for 16 weeks. (**C**) Representative image of eWAT isolated from WT and *LKO* mice fed a HFD for 16 weeks. (**D**) Quantification of Kupffer cells and monocytes. Hepatocytes were isolated from the *LKO* and WT mice and infiltration of monocytes and Kupffer cells were determined via FACS as explained in method section. All data represent mean ± SEM. *p<0.05 comparing *LKO* with WT mice using the unpaired t-test.

**Figure S3. The depletion of ANGPTL4 in the liver improves metabolic parameters without a change in food intake under WD fed conditions.** (**A**) Food consumption/day in WT and *LKO* mice fed WD for 16 weeks. (**B**-**D**) The fat mass measured by Echo-MRI, Plasma HDL-C levels and Fasting blood glucose level in WT and *LKO* mice injected with PCSK9-AAV and fed a WD for 16 weeks. (**E**) Representative cross-section analysis of macrophage content (CD68-positive cells) of the aortic root from WT and *LKO* mice fed a WD for 16 weeks. Quantifications of the graphs on the right. (**F**) Flow cytometry analysis of circulating monocytes, B cells, and T cells from WT and *LKO* mice fed a WD for 16 weeks. All data represent mean ± SEM. *p<0.05 comparing *LKO* with WT mice using the unpaired t-test.

**Figure S4. Genetic loss of ANGPTL4 in the liver induces fatty acid oxidation pathway and inhibits lipid biogenesis in HFD and WD fed mice.** (**A**-**D**) mRNA expression of PGC-1$\alpha$, CPT1A, HADHB, and ACC in the liver isolated from WT and *LKO* mice fed an HFD for 16 weeks. (**E**) Western blot showing p-ACC and ACC levels in the liver isolated from fasted WT and *LKO* mice fed an HFD for 16 weeks. The right panel shows p-ACC/ACC and ACC/HSP90 ratios from immunoblot images quantification. (**F** and **G**) mRNA levels of FASN and HMGCR in the liver from WT and *LKO* mice fed an HFD for 16 weeks. (**H-L**) mRNA levels of PGC-1$\alpha$, CPT1A, HADHB, ACC, and FASN in the liver of WT and *LKO* mice fed a WD for 16 weeks. R.E and R.D represents relative expression and relative density respectively. All data represent mean ± SEM. *p<0.05 **p<0.01 comparing *LKO* with WT mice using the unpaired t-test.

**Figure S5. GalNac-conjugated ANGPTL4 ASO treatment improves metabolic functions of mice.** (**A**) Body weight (n=5). (**B** and **C**) Lipoprotein profile of pooled plasma isolated from ANGPTL4 ASO or Ctrl ASO treated mice fed a CD (n=5). (**D**) Mean daily food intake in the HFD fed mice following 6-week injection of ANGPTL4 ASO or Ctrl ASO.

(**E**) Circulating blood counts from ANGPTL4 ASO or Ctrl ASO treated mice fed a HFD using Hemavet hematology analyzer. (**F** and **G**) Content of FPLC fractionated lipoproteins from pooled plasma of ANGPTL4 ASO or Cltr ASO injected mice fed a HFD (n=7). (**H**) Representative images of the liver, Liver weight and hepatic TAG levels from ANGPTL4 ASO or Ctrl ASO treated mice fed a HFD. All data represent mean ± SEM. *$p<0.05$ comparing ANGPTL4 ASO with Ctrl ASO treated mice using the unpaired t-test.

Figure S1

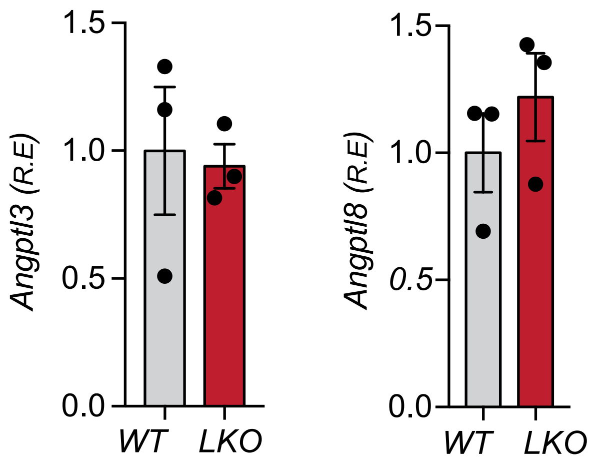

Figure S2

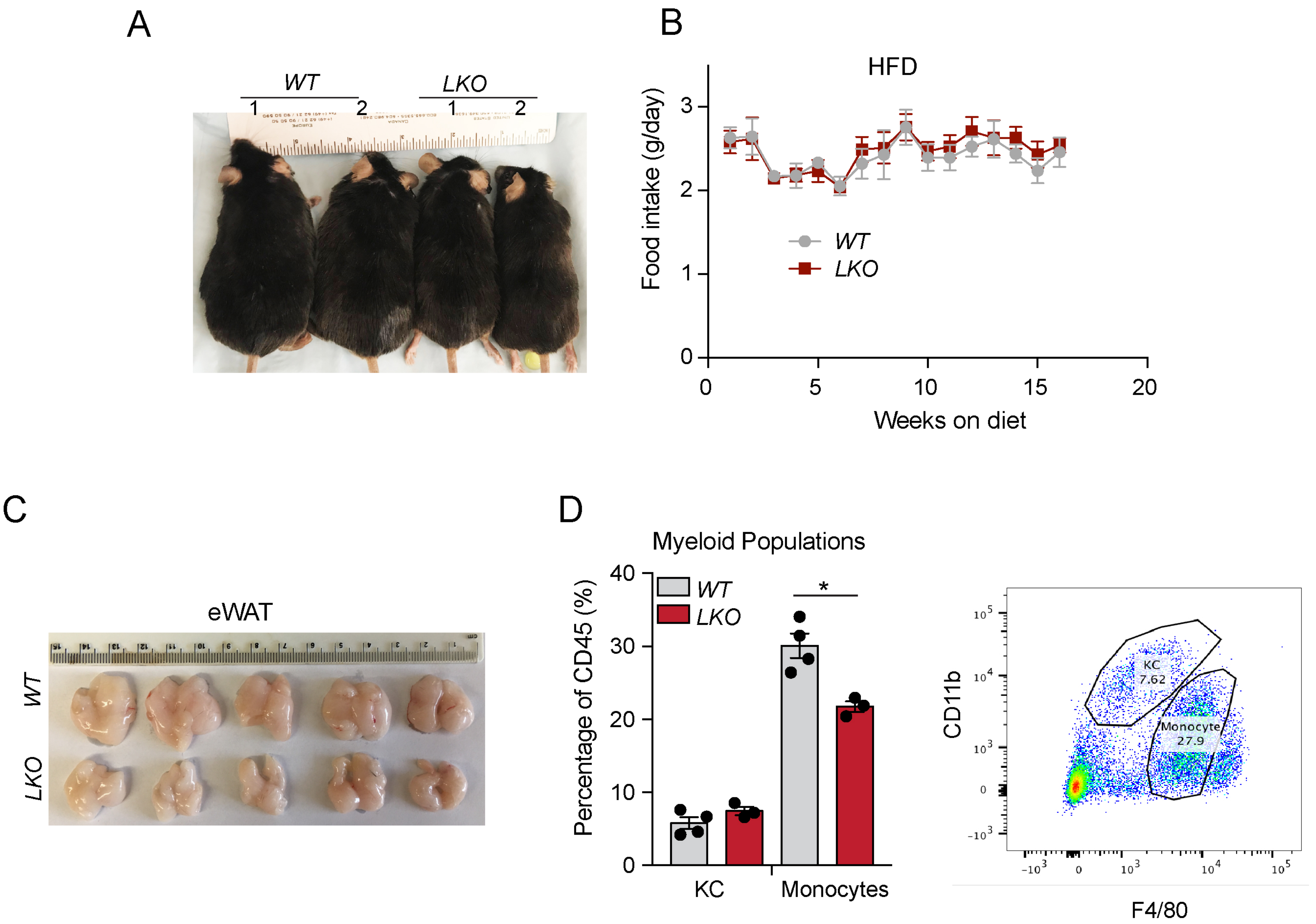
A
WT
LKO
1
2
1
2
B
HFD
Food intake (g/day)
Weeks on diet
WT
LKO
C
eWAT
WT
LKO
D
Myeloid Populations
Percentage of CD45 (%)
WT
LKO
*
KC
Monocytes
KC
7.62
Monocyte
27.9
CD11b
F4/80

### Figure S3

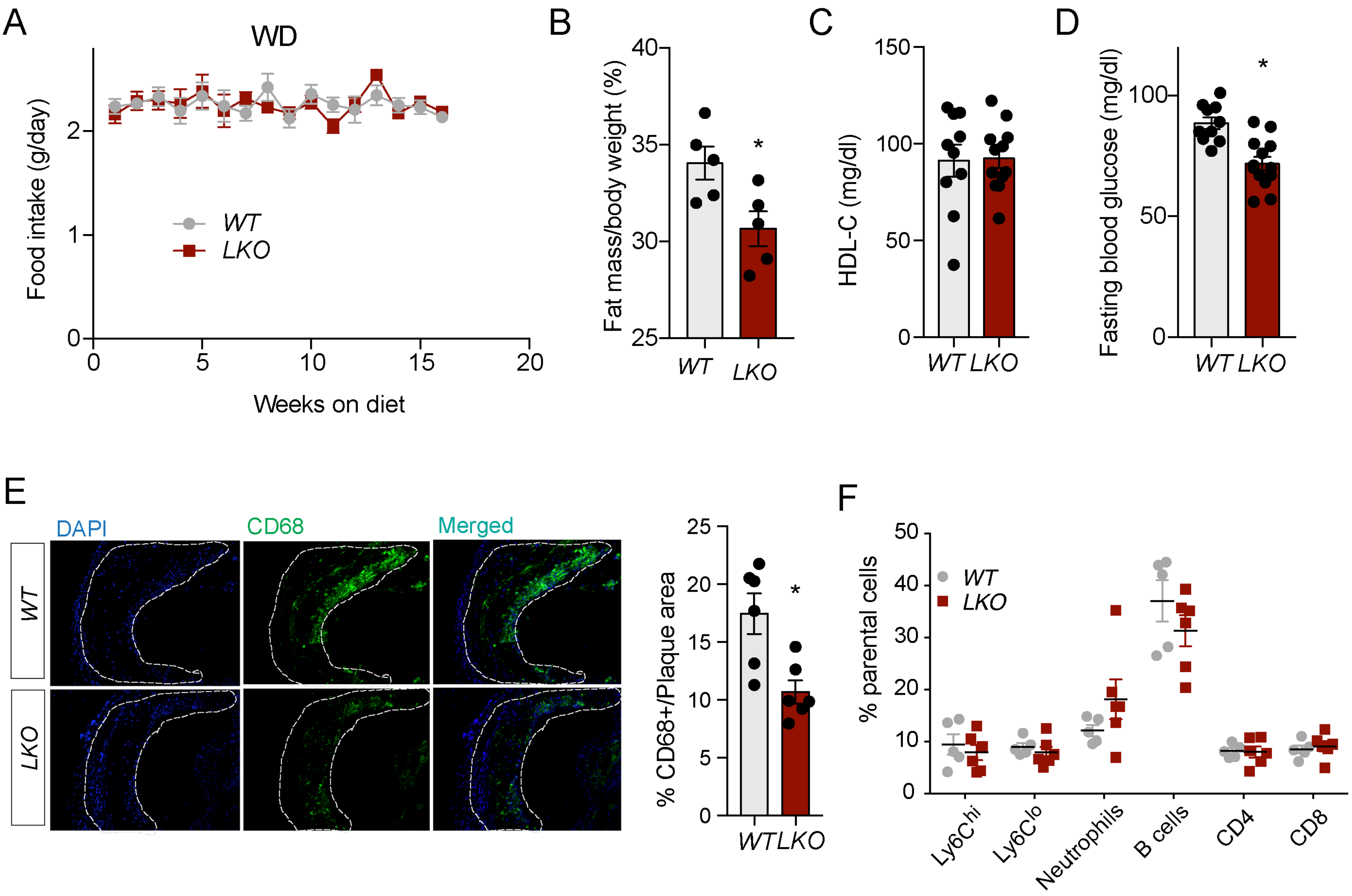

A
WD
Food intake (g/day)
Weeks on diet
WT
LKO
B
Fat mass/body weight (%)
C
HDL-C (mg/dl)
D
Fasting blood glucose (mg/dl)
E
DAPI
CD68
Merged
% CD68+/Plaque area
F
% parental cells
Ly6Chi
Ly6Clo
Neutrophils
B cells
CD4
CD8

### Figure S4

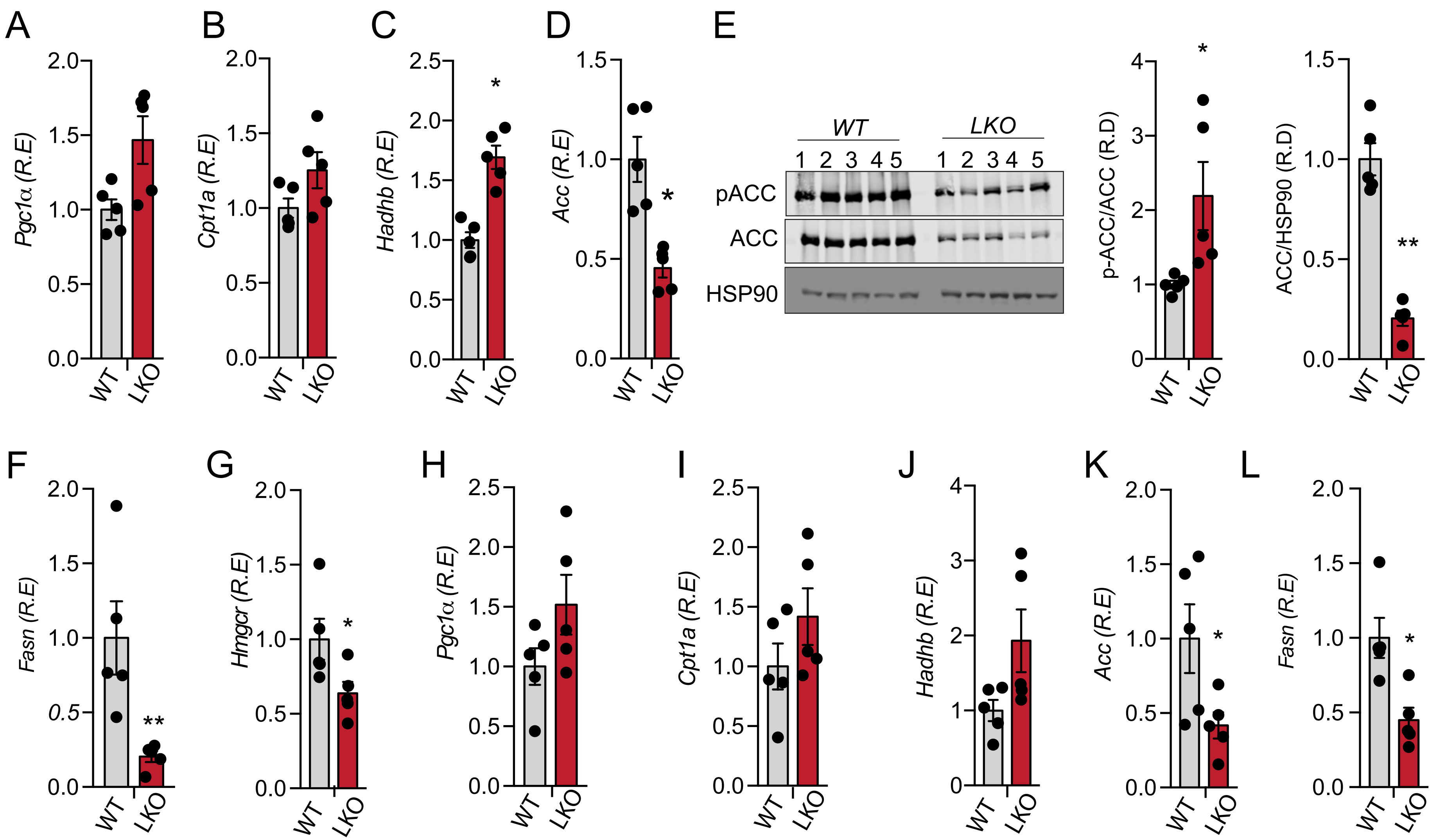
A
B
C
D
E
F
G
H
I
J
K
L
Pgc1α (R.E)
Cpt1a (R.E)
Hadhb (R.E)
Acc (R.E)
Fasn (R.E)
Hmgcr (R.E)
p-ACC/ACC (R.D)
ACC/HSP90 (R.D)
WT
LKO
1 2 3 4 5
pACC
ACC
HSP90
*
**

### Figure S5

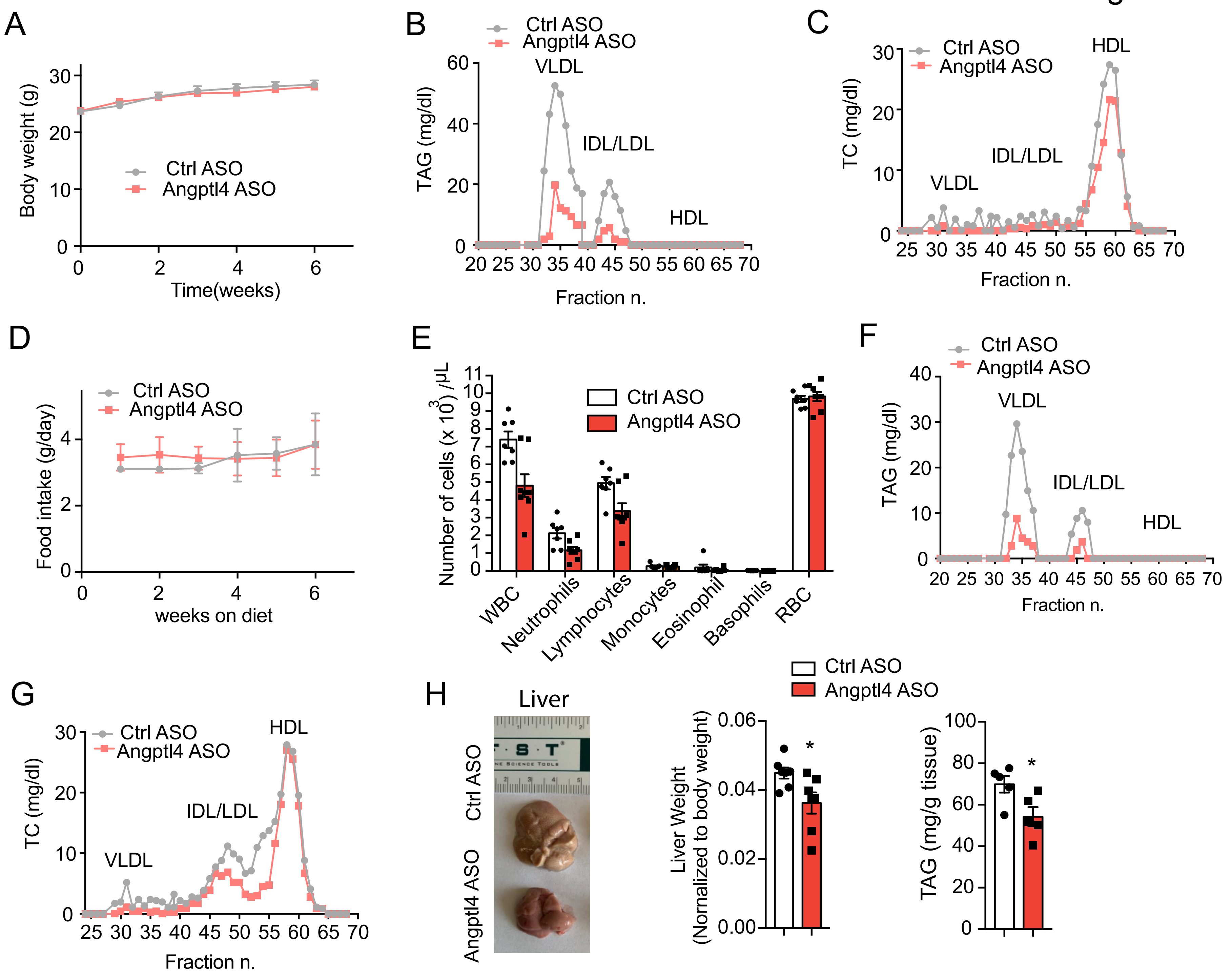
A
Body weight (g)
Time(weeks)
Ctrl ASO
Angptl4 ASO
B
TAG (mg/dl)
VLDL
IDL/LDL
HDL
Fraction n.
C
TC (mg/dl)
D
Food intake (g/day)
weeks on diet
E
Number of cells (x 10^3) /µL
WBC
Neutrophils
Lymphocytes
Monocytes
Eosinophil
Basophils
RBC
F
G
H
Liver
Ctrl ASO
Angptl4 ASO
Liver Weight
(Normalized to body weight)
TAG (mg/g tissue)
*
